# Hydrostatic pressure shapes and canalizes semicircular canal morphology to ensure vestibular function

**DOI:** 10.64898/2026.07.22.740136

**Authors:** Jiacheng Wang, Kira L. Heikes, Yufei Wu, Anne Belle Briggs, Kevin Li, Nadia Eliora Suriato, Srihas Surapaneni, Roarke Horstmeyer, Michel Bagnat, Akankshi Munjal

## Abstract

How tissues acquire reproducible shapes to support their function is a fundamental question in developmental biology. Zebrafish semicircular canals form when epithelial pillars partition the lumen of the otic vesicle, the embryonic precursor of the inner ear, into the tubes that sense head rotation for balance. We show that hydrostatic pressure generated by the inflating otic vesicle shapes pillar geometry. Acute vesicle deflation shortens and broadens pillars, whereas pharmacologically induced inflation elongates and narrows them, effects captured by a physical model that treats the vesicle as a pressurized viscoelastic shell. Pillars initially form with variable curvature, but continued vesicle expansion drives them toward a common, straight geometry, a process accelerated by inflation and blocked by deflation. We identify Wnt signaling as crucial for sustaining this pressure by maintaining epithelial barrier integrity. Its disruption impairs pillar geometry and vestibular-dependent swimming behavior. Together, these findings identify hydrostatic pressure as a critical physical mechanism that canalizes tissue geometry to ensure robust organ formation and function.

## Introduction

Organ function is tightly linked to organ geometry. In the heart, ventricular curvature determines pumping efficiency^1^; in the lung, the alveolar surface-area-to-volume ratio constrains gas exchange^2–4^; and in the gut, villus height and spacing regulate nutrient absorption^5,6^. Among the most striking examples of form-function coupling are the evolutionarily conserved semicircular canals of the vestibular system in the inner ear, which detect head rotation through the motion of a fluid called endolymph within these curved, mutually orthogonal tubes^7,8^. Analyses of semicircular canal structure and function predict that vestibular sensitivity depends strongly on canal geometry^9,10^. According to Poiseuille’s law, small changes in canal radius or length would substantially alter endolymph flow, rendering vestibular function highly sensitive to minor geometric deviations. How such precise organ geometry is produced reproducibly and robustly during development remains a fundamental open question in biology.

Inner ear development occurs within the otic vesicle, a fluid-filled epithelial cyst that arises from the otic placode, a thickened region of cranial ectoderm adjacent to the hindbrain^11^. In zebrafish, the placode undergoes cavitation driven by ion transport, generating an osmotic gradient that draws fluid into the nascent lumen to form the otic vesicle^12,13^. This establishes a pressurized lumen enclosed by a tight epithelial barrier. Between ∼16 and 45 hpf, continued fluid accumulation increases luminal hydrostatic pressure and drives vesicle expansion^13^.

Semicircular canal formation begins at ∼45 hours post-fertilization (hpf), when six epithelial buds extend into the lumen through osmotic swelling of a hyaluronan-rich extracellular matrix (HA-ECM)^14,15^. Adjacent buds then contact and fuse to form three transluminal pillars that partition the lumen into semicircular canals, coincident with the cessation of HA-ECM synthesis^14,16–18^. Mature semicircular canals detect angular acceleration through head rotation–driven flow of the luminal fluid, termed endolymph, which deflects sensory hair cells at the base of each canal^19^. Canal geometry is therefore defined by the relationship between pillar shape and the surrounding epithelial walls that form the canal tubes. How this relationship is established during morphogenesis to generate stereotyped canal architecture remains unclear.

Using the optical and physical accessibility of the zebrafish otic vesicle, combined with perturbation approaches, theoretical modeling, and behavioral assays, we investigated how vesicle expansion influences semicircular canal morphology. We find that pressure-driven inflation of the otic vesicle plays a central role in shaping canal architecture. Newly formed pillars remain mechanically coupled to the pressurized lumen: acute deflation shortens and widens pillars, whereas pharmacologically induced expansion produces longer, narrower pillars, consistent with predictions from a physical model of a pressurized epithelial shell. Although pillars initially form with substantial geometric variability, continued vesicle expansion progressively straightens them, canalizing pillar morphology. Pharmacologically induced inflation is sufficient to accelerate pillar straightening, whereas acute otic vesicle deflation causes pillar bending, demonstrating that luminal hydrostatic pressure is persistently required to establish and maintain pillar geometry. Finally, we show that Wnt signaling regulates pillar morphology by controlling vesicle expansion through epithelial barrier function; its inhibition causes vesicle deflation, disrupts pillar geometry, and leads to vestibular dysfunction in larvae. Together, these results identify hydrostatic pressure-driven lumen expansion as a key mechanical regulator of both pillar morphology and the canalization of pillar geometry, thereby ensuring robust semicircular canal architecture and vestibular function.

## Results

### Hydrostatic pressure continues to drive otic vesicle expansion during semicircular canal morphogenesis

Previous work has shown that during early otic vesicle growth (16–45 hpf), fluid accumulation drives luminal expansion that outpaces tissue growth, leading to a progressive increase in hydrostatic pressure^13,20^. Consistent with this, reanalysis of published volumetric data indicates that the tissue fraction of the otic vesicle defined as tissue volume/total volume, decreases from ∼67% at 30 hpf to ∼44% at 45 hpf, reflecting a transition from a tissue-dominated to a lumen-dominated structure (Supp Fig. 1). During this window, pressure increases from ∼110 Pa to ∼215 Pa, with the epithelium thinning under this increasing pressure^13^. These observations support the use of tissue fraction as a proxy for the pressurized state of the otic vesicle. Whether this relationship persists during later stages of inner ear development remains unclear. If this relationship persists, sustained pressure may continue to play a mechanical role during semicircular canal morphogenesis, which begins at ∼45 hpf as the vesicle reaches a critical size.

**Figure 1.**
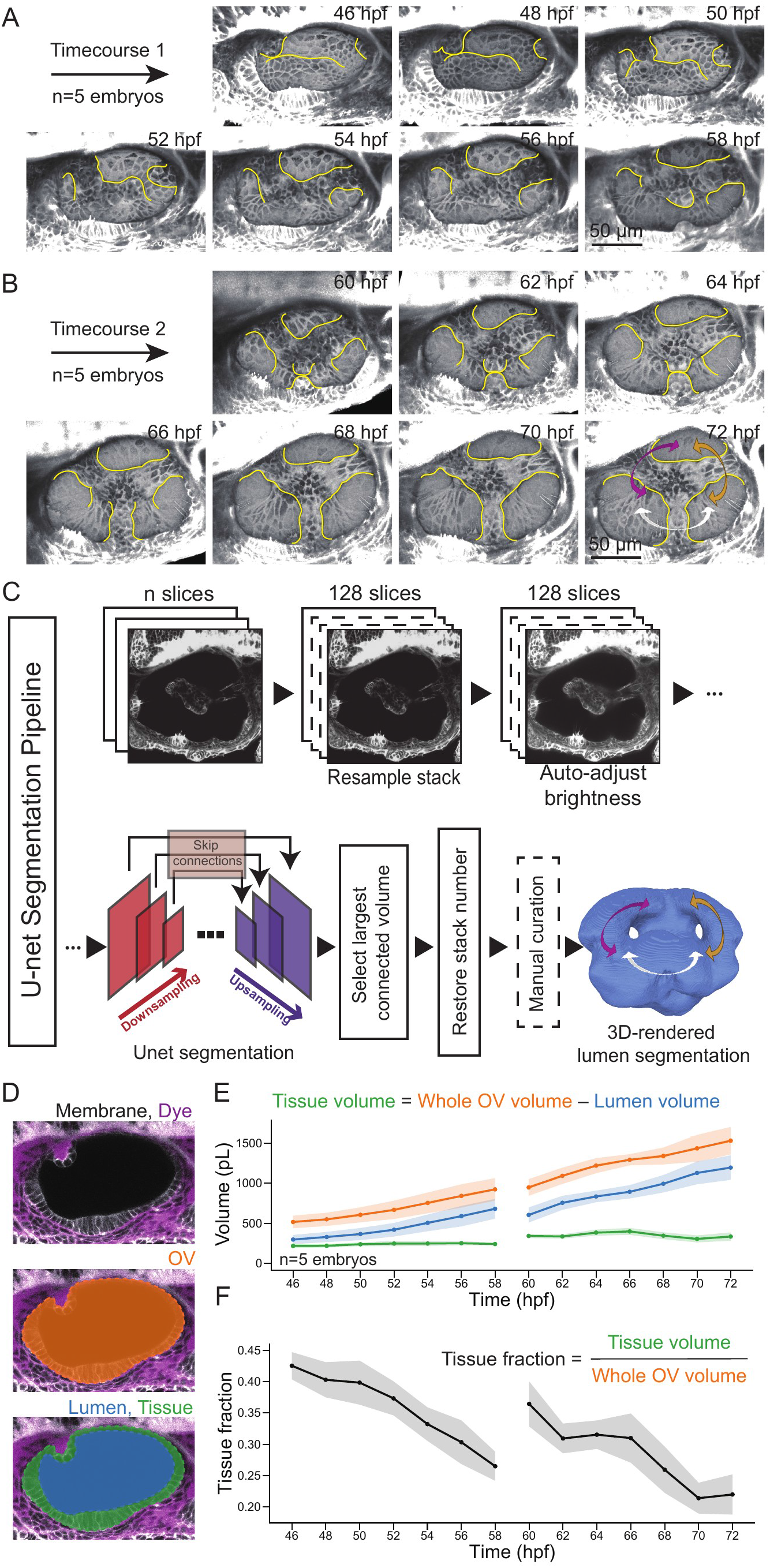
Hydrostatic pressure continues to drive otic vesicle expansion during semicircular canal morphogenesis. A-B. Representative images of the timecourse from 46 to 58 hpf (A) and from 60 to 72 hpf (B). Bud and pillar regions are outlined in yellow. Semicircular canals are denoted in the 3D-rendered 72 hpf image by arrows (anterior canal, purple; posterior canal, brown; lateral canal, white). **C.** Volume measurement pipeline, including image preprocessing, a trained UNet structure for segmenting the otic vesicle (OV) lumen space, and occasional manual curation in ITK-Snap. Semicircular canals are denoted in the 3D-rendered lumen segmentation by arrows (anterior canal, purple; posterior canal, brown; lateral canal, white). **D.** Image overlay defining the volumes plotted in E: Whole OV in orange, lumen in blue, and tissue layer in green. **E.** Volumes measured by the Unet pipeline over time for the timecourses from 46 to 58 hpf and from 60 to 72 hpf. Data points indicate the mean, and line thickness represents the standard deviation from the mean. **F.** Tissue fraction as a function of tissue volume divided by whole OV volume over time for the timecourses. Scale bars = 50 µm. n=5 embryos per timecourse.

To test this, we quantified vesicle growth during semicircular canal formation across two developmental windows (46–58 hpf and 60–72 hpf), spanning key remodeling events associated with canal morphogenesis: sequential bud formation and their fusion to give rise to three pillars (Fig. 1A, B). To accurately capture developmental dynamics, embryos in timecourse experiments were mounted only briefly for imaging and returned to normal conditions between time points, to avoid the developmental delay associated with continuous timelapse imaging under a coverslip. To extract lumen and tissue volumes from these 3D datasets, we developed a U-Net^21^–based segmentation pipeline (Fig. 1C) that enabled automated, high-throughput segmentation of the lumen and otic epithelium across all morphogenetic stages (Supp. Video 1). Lumen volume was calculated by segmenting the fluid-filled cavity, and tissue volume was calculated by subtracting the lumen volume from the total otic vesicle volume, as defined by the epithelial boundary (Fig. 1D).

We found that lumen volume continues to increase approximately linearly during canal morphogenesis, whereas tissue volume remains largely unchanged (or decreases slightly) between 46–72 hpf (Fig. 1E). As a result, the tissue fraction progressively decreases over this period, from ∼42% to ∼22%, indicating that the otic vesicle becomes increasingly fluid-dominated (Fig. 1F). The continued increase in lumen volume in the absence of significant tissue growth, reflected by the decreasing tissue fraction, supports a model in which otic vesicle inflation during canal morphogenesis is primarily driven by fluid accumulation generating sustained hydrostatic pressure. We therefore asked whether this sustained pressure mechanically shapes pillar formation.

### Otic vesicle expansion and early pillar morphology are mechanically coupled

Semicircular canal morphogenesis begins with the extension of epithelial buds and culminates in their fusion to form three transluminal pillars that partition the otic vesicle into three canals^12^. To test whether hydrostatic pressure is required for this early phase of bud growth, previous studies ablated the vesicle with a two-photon laser^14^, an approach that rapidly relieves luminal pressure by disrupting the epithelial barrier that maintains it^13^. Deflation of the otic vesicle has little effect on bud extension, which is primarily driven by osmotic swelling of the HA-ECM beneath the budding epithelium^14^. However, whether otic vesicle expansion contributes to later stages of morphogenesis, including pillar formation and shaping, has not been examined.

To address this, we performed laser ablation of the OV immediately after pillar formation, resulting in rapid vesicle deflation, with lumen volume decreasing by an average of ∼29% within 30 minutes post-ablation (Fig. 2A-C, Supp Fig. 2A-B’’’). This perturbation led to a marked geometric transformation of newly formed pillars, which became shorter and wider with a significant reduction in pillar aspect ratio (length/width) compared to pre-ablation (Fig. 2D). Conversely, increasing vesicle expansion with forskolin, a cAMP agonist that stimulates ion transport and thereby osmotic fluid influx into the lumen^22^, increased lumen volume by an average of ∼43% over 4 hours. This had the opposite effect, producing longer, narrower pillars and a correspondingly increased pillar aspect ratio (Fig. 2E-J, Supp Fig. 2C-F’’’). Together, these results suggest that pillar geometry is mechanically coupled to the inflation state of the otic vesicle lumen.

**Figure 2.**
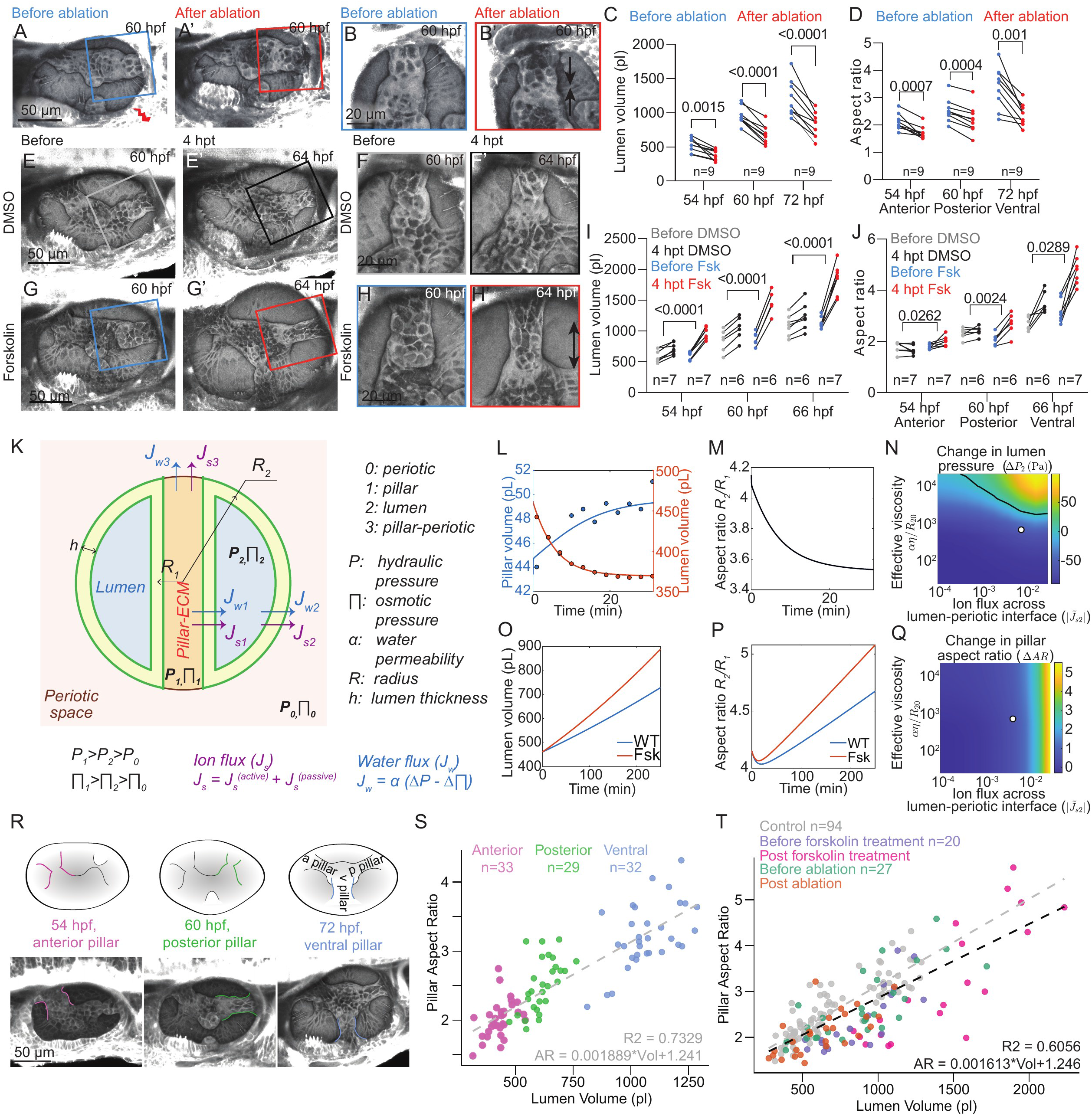
Otic vesicle expansion and early pillar morphology are mechanically coupled. A-A’. Representative images of an OV at 54 hpf before and after ablation. **B-B’.** Magnified insets of the boxed regions in A-A’ showing relaxation of the pillar and decreased aspect ratio. **C.** Quantification of OV lumen volume before and after ablation at the three developmental stages indicated. P-values as labeled (two-tailed paired t-test). **D.** Quantification of pillar aspect ratio before and after ablation at the developmental stages indicated. Pillar measured at each staged ablation is indicated along the x-axis. P-values as labeled (two-tailed paired t-test). **E-E’**. Representative images of an OV before and 4 hours post-treatment (hpt) by soaking in DMSO. **F-F’**. Magnified insets of the boxed regions in E-E’ showing minimal change in pillar aspect ratio after DMSO treatment. **G-G’.** Representative images of an OV before and 4 hpt by soaking in Forskolin. **H-H’.** Magnified insets of the boxed regions in G-G’ showing elongation of the pillar and increase in aspect ratio after Forskolin treatment. **I.** Quantification of OV lumen volume before and 4 hpt in DMSO or Forskolin at the developmental stages indicated. **J.** Quantification of pillar aspect ratio before and 4 hpt in DMSO or Forskolin at the developmental stages indicated. Pillar measured at each staged treatment is indicated along the x-axis. **K.** Schematic of the fluid driven physical model of OV lumen and pillar dynamics. Symbols are defined to the right of the schematic, along with assumptions made in the model, and the equations of water and ion flux across boundaries. **L.** Computed values of pillar volume (blue, pL) and lumen volume (orange, pL) over time after simulated OV ablation. Scatter points indicate experimentally measured values. **M.** Computed values of effective pillar aspect ratio over time after simulated OV ablation. **N.** Computed phase diagram of change in lumen pressure (kPa) over a four-hour period as a function of lumen ion influx and effective epithelial viscosity. **O.** Computed values of lumen volume (pL) over time for wild type (WT, blue) and Fsk (red) conditions. **P.** Computed values of effective pillar aspect ratio over time defined as a function of lumen radius divided by pillar radius for WT (blue) and Fsk (red) conditions. **Q.** Computed phase diagram showing the change in aspect ratio (Δ *AR*) over a four-hour period as a function of lumen ion influx (|*J*^∼^_s2_|) and epithelial viscosity (*αη*/*R*_2O_). Here *R*_2O_ denotes the reference lumen radius used for nondimensionalization. **R.** Representative images of OVs in the staged time series used for the quantification in S. **S.** Quantification of pillar aspect ratio and OV lumen volume (pL) for embryos pooled at the following developmental stages: anterior pillars (pink) quantified at 54 hpf, posterior (green) pillars measured at 60 hpf, and ventral (blue) pillars measured at 72 hpf. Linear fit to the data shown as dashed line. **T**. Quantification of posterior pillar aspect ratio and OV lumen volume (pL) for control embryos from S (grey), before and after soaking with Forskolin (purple and magenta), and before and after OV ablation (green and orange). Linear fit to the control data shown as grey dashed line. Linear fit to all data shown as black dashed line. “n” denotes number of embryos. Scale bar values indicated in each image. P-values as labeled (two-tailed paired t-test).

To understand how this coupling arises, we developed a fluid-driven physical model that treats the otic vesicle as a spherical viscoelastic epithelial shell (radius *R*₂) enclosing an osmotically active lumen and a single internal pillar-like structure (Fig. 2K). The model comprises three compartments—the pillar (1), lumen (2), and periotic space (representing the space surrounding the otic vesicle) (0)—connected by pillar–lumen, lumen–periotic, and pillar–periotic interfaces. Ion flux (*J*_s_) combines passive diffusion and active transport. Consistent with previous theoretical frameworks for epithelial hydraulics and lumen growth^23–25^, this flux creates osmotic gradients to drive water accumulation in the lumen, giving rise to hydrostatic pressure that resists further influx. The water flux (*J*_w_) is therefore driven by osmotic and hydrostatic pressure differences across each interface. The lumen and pillar walls are modeled as thin viscoelastic epithelial layers of thickness *h*, in which elasticity (*E*) determines the magnitude of stresses generated by deformation, whereas viscosity (*η*) determines how rapidly these stresses relax. Although HA synthesis ceases after bud fusion^14,16–18^, we observed that residual HA remains within pillars (Supp. Fig. 2G). The pillar was therefore modeled as a triphasic hydrogel, whose mechanics arise from osmotic swelling and elastic deformation^26^. Even where the pillar opens into the periotic space, the gel matrix defines the pillar–periotic interface (Supp. Fig. 2H). Hydrostatic pressure in the lumen and mechanical stress within the pillar are balanced by epithelial tension through the Young–Laplace relation. Together, these properties couple tissue mechanics to hydrostatic pressure. Full derivations, nondimensionalization, parameters, and numerical implementation are provided in the Methods Theoretical Modeling section. Using experimentally measured vesicle and pillar dimensions together with literature-derived material parameters (Methods Theoretical Modeling section), the model converged to a steady state in which the pillar is the most pressurized compartment, followed by the lumen and periotic space (Fig. 2K).

The model captures both short-timescale responses to acute perturbations and long-timescale growth dynamics. To model pillar response during otic vesicle deflation after ablation, water movement was described as hydrostatic pressure-driven flow through a puncture in the lumen wall (Methods Theoretical Modeling section). Because ablation transiently perturbs mechanical equilibrium, volume relaxation reflects epithelial wall viscosity. We therefore measured the dynamics of lumen and pillar volumes post-ablation to constrain lumen and pillar wall viscosities (Fig. 2L). A literature-reported otic epithelium viscosity^13^ accurately reproduced the lumen volume dynamics (Fig. 2L), whereas the pillar wall viscosity was fit to the measured pillar volume dynamics, yielding a lower effective viscosity (Methods Theoretical Modeling section and Fig. 2L). Using these viscosity values, the model accurately reproduced the experimentally observed decrease in pillar aspect ratio after ablation (Fig. 2M). Sensitivity analysis further predicted that the two wall viscosities make distinct contributions: lumen wall viscosity primarily governs lumen volume relaxation, whereas pillar wall viscosity primarily governs pillar volume and geometry (Supp Fig. 2I-K). Together, these results show that, within a physiological range of wall viscosities, the model recapitulates changes in pillar aspect ratio in response to acute deformation and suggests that the pillar wall responds more rapidly to mechanical deformation than the lumen wall, likely due to its faster viscoelastic stress relaxation.

To model long-timescale growth, we assumed luminal fluid accumulation via active ion influx with no tissue growth, consistent with the measured increase in lumen volume and decrease in tissue fraction (Fig. 1E-F and Methods Theoretical Modeling section). Varying tissue viscosity and ion flux revealed that luminal pressure depends jointly on both parameters, with a critical boundary separating regimes of increasing and decreasing luminal pressure (Fig. 2N). Using the literature-derived viscosity value as above^13^, the model predicts that luminal pressure gradually decreases due to viscous relaxation before stabilizing (Fig. 2N, white circle). Because tissue viscosity may itself change during development^27^, and both tissue viscosity and luminal pressure dynamics are challenging to measure accurately *in vivo*, our current data cannot distinguish whether luminal pressure gradually declines, is maintained, or even rises before changes in tissue material properties become significant. Nevertheless, across the physiological range of wall viscosities examined, the model consistently predicts that ion-flux-driven fluid accumulation elongates and narrows the pillar, and that increasing ion influx further enhances lumen expansion and pillar aspect ratio, accurately recapitulating the effects of forskolin treatment (Fig. 2O-Q).

While the pillar aspect ratio is largely insensitive to tissue viscosity, it depends strongly on ion flux (Fig. 2Q), suggesting that the resulting hydrostatic pressure, and therefore the inflation state of the otic vesicle, determines pillar geometry. Because the anterior, posterior, and ventral pillars form sequentially (Fig. 1B–C), developmental time and vesicle inflation are naturally correlated. The model therefore predicts that, despite this correlation, the geometry of newly formed pillars should reflect the inflation state of the vesicle rather than developmental time. Because pillars flatten and acquire more complex geometry as they mature, we could not track aspect ratio over time within individual pillars. Instead, we quantified the geometry of newly formed pillars across a large cohort of embryos spanning the full range of otic vesicle volumes observed during pillar morphogenesis (54–72 hpf) (Fig. 2R). Pillar aspect ratio scaled linearly with otic vesicle volume (Fig. 2S; R² = 0.732), with newly formed anterior, posterior, and ventral pillars exhibiting progressively greater aspect ratios as they formed at increasingly inflated stages. Within each pillar class, aspect ratio likewise scaled with otic vesicle volume, following the same volume-dependent relationship (Fig. 2S). Importantly, this relationship was preserved in embryos subjected to either inflation or deflation perturbations (Fig. 2T, R² = 0.605). Together, these findings demonstrate that the geometry of newly formed pillars is mechanically coupled to otic vesicle inflation, a relationship quantitatively explained by a fluid-driven physical model that correctly predicts pillar geometry across both acute perturbations and developmental growth.

### Otic vesicle expansion canalizes variability in pillar morphology

Having established that the inflation state determines the initial geometry of pillars, we next asked how pillar morphology evolves during continued otic vesicle expansion. We observed that opposing buds fuse at variable angles (Fig. 3A and Fig. 3D, green), resulting in newly formed pillars with substantial variability in curvature (Fig. 3B, Supp. Fig. 3A, B). To quantify this variability, we measured pillar straightness in three dimensions as the angle between the two fused bud arms, calculated as the arccosine of the dot product of their normalized direction vectors (Fig. 3C). Across the same large dataset used to measure pillar aspect ratio (Fig. 2S), pillar angles increased progressively toward 180° with increasing otic vesicle volume (Fig. 3D, Supp. Fig. 3C, D), while the coefficient of variation in pillar angle decreased (Fig. 3E, Supp. Fig. 3E, F), indicating that initially heterogeneous pillar geometries progressively converged toward a common straight morphology. Thus, pillar shape becomes progressively canalized during development.

**Figure 3.**
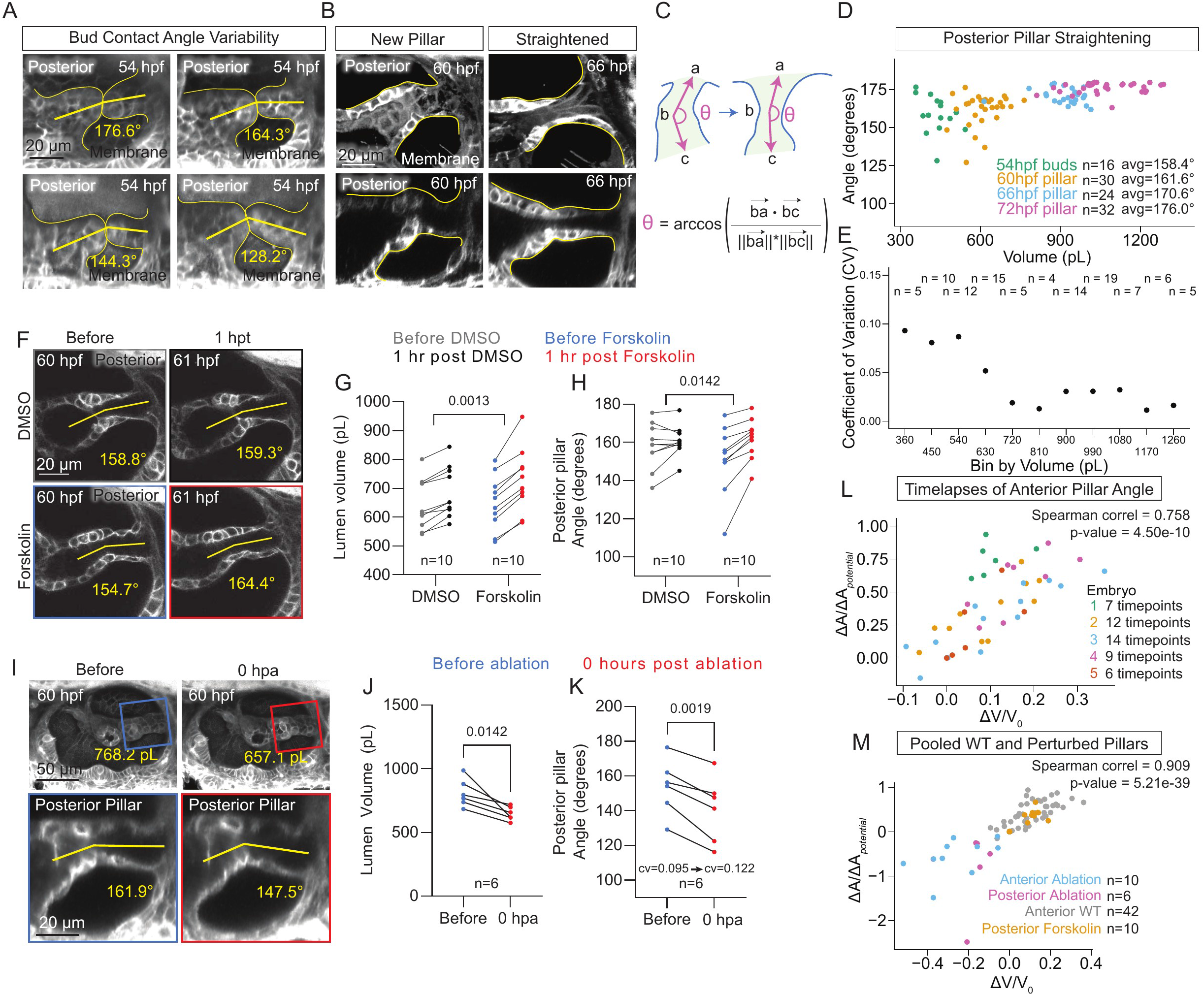
Otic vesicle expansion canalizes variability in pillar morphology. **A.** Representative confocal images of 3D-rendered posterior buds at contact from membrane mNeonGreen-expressing embryos at 54 hpf, demonstrating initial variability in bud contact angle. **B.** Representative confocal images of 3D-rendered posterior pillars from membrane-mNeonGreen-expressing embryos at the stages indicated, demonstrating straightening of posterior pillars after initial shape variability. **C.** Schematic defining pillar angle metric used to describe pillar straightness quantitatively. **D.** Scatter plot of canalization of posterior bud contact or pillar angle (degrees) vs otic vesicle lumen volume (pL) quantified in embryos pooled at the following developmental stages: 54 hpf (green, bud contact), 60 hpf (orange, pillars), 66 hpf (blue, pillars), and 72 hpf (pink, pillars). **E.** Plot of coefficient of variation calculated from embryos in D binned by lumen volume in 90-pL bins. **F.** Representative confocal images of 3D-rendered posterior pillars from membrane-mNeonGreen-expressing embryos at the stages indicated, before and 1 hour post treatment (hpt) with DMSO as a control or Forskolin in DMSO. **G.** Quantification of OV lumen volume (pL) before and 1 hpt in DMSO or Forskolin with DMSO at 60 hpf. P-values as labeled (two-tailed unpaired t-test). **H.** Quantification of posterior pillar angle (degrees) before and 1 hpt in DMSO or Forskolin with DMSO at 60 hpf. P-values as labeled (two-tailed unpaired t-test). **I.** Representative confocal images of 3D-rendered OVs from membrane-mNeonGreen-expressing embryos at the stages indicated, before and after 2-photon laser ablation of the OV to reduce lumen volume and pressure. **J.** Quantification of OV lumen volume (pL) before and after 2-photon laser ablation of the OV at 60 hpf. P-values as labeled (two-tailed paired t-test). **K.** Quantification of posterior pillar angle (degrees) before and after 2-photon laser ablation of the OV at 60 hpf. P-values as labeled (two-tailed paired t-test). **L.** Plot of change in anterior pillar angle (degrees, A_n_-A_0_) scaled to initial potential for pillar straightening at pillar formation (180°-A_0_) vs change in OV lumen volume (pL, V_n_-V_0_) scaled to initial OV lumen volume at pillar formation (V_0_); data pooled from all timepoints (n=42) quantified from 5 embryos that were timelapse imaged from pillar formation to pillar straightening (colored by embryo). Spearman coefficient of pooled data as shown (0.758). **M.** Plot of scaled change in pillar angle (degrees) vs scaled change in OV lumen volume (pL); data pooled from all timepoints quantified from 5 time-lapsed embryos (anterior pillar in wild type, WT, grey) and for perturbed conditions (anterior pillar from 54 hpf ablation, blue, posterior pillar from 60hpf Ablation, pink, posterior pillar from 60 hpf Forskolin, orange). Change quantified from before and after perturbation. Spearman coefficient of pooled data as shown (0.909). Coefficient of variation (cv) indicated in certain graphs. Scale bar values indicated in each image. “n” denotes number of embryos.

We next investigated the mechanism underlying pillar straightening. We first hypothesized that residual swelling pressure within the HA-ECM extracellular matrix continues to mechanically influence pillar geometry after fusion^14^. To test this hypothesis, we enzymatically degraded HA by injecting hyaluronidase (HAase) into the periotic space, thereby eliminating its associated swelling pressure^14^. Surprisingly, HA digestion did not impair pillar straightening but instead significantly accelerated it relative to PBS-injected controls (Supp. Fig. 3G, H). These findings indicate that the residual HA-ECM matrix transiently resists, rather than drives, pillar straightening, thereby slowing the canalization of pillar morphology.

We therefore asked whether continued otic vesicle inflation provides the mechanical input that remodels pillar geometry after fusion. In the model, ion-flux-driven luminal hydrostatic pressure inflates the otic vesicle, thereby generating mechanical stresses within the surrounding epithelium, including the pillar walls (Fig. 2K). We therefore perturbed luminal pressure *in vivo* and measured consequent change in pillar straightness. Increasing luminal pressure with forskolin accelerated otic vesicle inflation and induced rapid pillar straightening within one hour, demonstrating that increased inflation is sufficient to accelerate this process (Fig. 3F-H). Conversely, reducing luminal pressure by targeted laser ablation rapidly deflated the otic vesicle and immediately increased pillar bending, demonstrating that continued luminal pressure is required to maintain pillar straightness (Fig. 3I-K, Supp. Fig. 3I-K). Together, these acute perturbations demonstrate that pillar geometry responds rapidly to changes in pressure-driven inflation of the otic vesicle.

If continued otic vesicle inflation drives pillar straightening, then the rate of straightening should be determined by the rate of inflation of the otic vesicle rather than developmental time itself. To test this prediction, we tracked individual pillars by time-lapse imaging from their formation through straightening (Supp. Fig. 3L-P, Q). To account for differences in starting otic vesicle lumen volume and how much pillar straightening was still possible, we normalized volume change to each embryo’s initial lumen volume and angle change to the pillar’s remaining distance from straight. We observed a clear positive correlation between lumen inflation and pillar straightening in wild-type embryos (Fig. 3L, Spearman correlation = 0.758). Because each embryo samples only a relatively narrow range of inflation states, we next expanded the analysis by combining wild-type embryos with inflation-and deflation-perturbed embryos. Across this broader range of lumen volumes, the association between otic vesicle inflation and pillar straightening became even more pronounced (Fig. 3M, Spearman correlation = 0.909), supporting the idea that lumen inflation promotes pillar straightening. Together, these findings reveal that continued pressure-driven otic vesicle inflation progressively remodels initially heterogeneous pillar geometries, ultimately canalizing pillar morphology during semicircular canal morphogenesis.

### Wnt signaling regulates otic vesicle expansion by maintaining epithelial barrier function during canal morphogenesis

We next sought to identify biological regulators that control the continued expansion of the otic vesicle during semicircular canal morphogenesis. Because vesicle expansion depends on fluid accumulation within the lumen, we reasoned that signaling pathways regulating epithelial ion transport or epithelial barrier function could influence this process^24,28,29^. To identify candidate regulators, we examined publicly available single-cell transcriptomic datasets of the developing zebrafish inner ear. Analyses of both the embryo-wide Daniocell atlas^30^ (Fig. 4A-B) and an inner ear–specific single-cell dataset^31^ (Supp. Fig. 4A-E) revealed strong expression of multiple components of the Wnt signaling pathway in the non-sensory dorsal region of the otic vesicle that gives rise to canal structures^16^, including *wls* (*wntless*). *Wls* encodes a conserved transmembrane cargo receptor required for the secretion of Wnt ligands^32^ and is broadly expressed throughout the otic epithelium^33^. Using Hybridization Chain Reaction-Fluorescent In Situ Hybridization (HCR-FISH)^34^, we observed enriched expression of *wls* in the canal-forming regions (Fig. 4C). Consistent with a potential role in otic vesicle growth, previous work reported that *wls* mutants develop reduced otic vesicles at 4 days post fertilization^35^. We reasoned that this reduction in size may reflect mis-regulated vesicle inflation. Because Wls is required for the secretion of Wnt ligands, targeting *wls* provides a means to broadly disrupt Wnt signaling within the developing inner ear.

**Figure 4.**
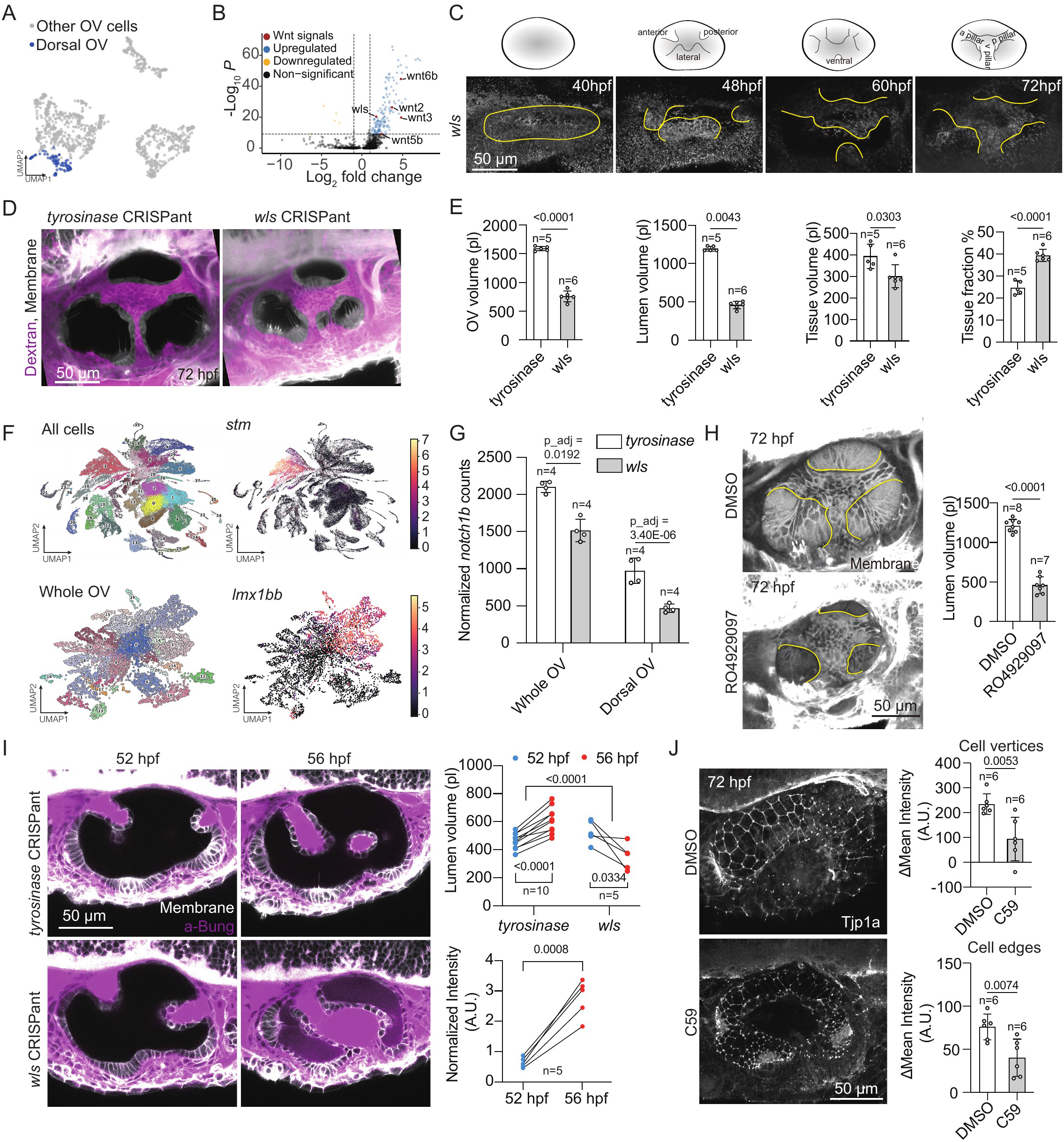
Wnt signaling regulates OV expansion by maintaining epithelial barrier function during canal morphogenesis. **A.** UMAP dimensionality reduction of OV cells in the published embryo-wide Daniocell scRNAseq atlas, with dorsal OV cells highlighted in blue, defined by counts of the transcription factor gene *LIM homeobox transcription factor 1, beta b* (*lmx1bb*). **B.** Volcano plot showing differential gene expression between dorsal OV cells and all other OV cells. Genes meeting the significance criteria (adjusted P < 10^-9^ and log2 fold change > 1, blue; log2 fold change < −1, yellow) are colored by direction of change; non-significant genes are shown in black. Several genes related to Wnt signaling are highlighted (red), including *wntless* (*wls*). **C.** Representative images of 3D-rendered OVs at the developmental stages indicated, stained with multiplex *in situ* probes against *wls*. Illustrations show typical progression of OV morphogenesis, highlighting the projections that fuse into the three pillars: anterior, posterior, and ventral. **D.** Representative images of OVs from membrane-mNeonGreen-expressing embryos at 72 hpf after Crispr-Cas9 knockdown of *tyrosinase* (control) or *wls*. Prior to imaging, embryos were injected periotically with Dextran conjugated with Texas Red to visualize the outer surface of the OV. **E.** Quantification of whole OV volume (pL), OV lumen volume (pL), epithelial tissue volume (pL), and tissue fraction (%), defined as the tissue volume divided by the whole OV volume in embryos at 72 hpf after Crispr-Cas9 knockdown of either *tyrosinase* (control, white) or *wls* (grey). P-values as labeled (Tissue fraction and OV volume: unpaired, two-tailed Student’s t-test; Tissue volume and lumen volume: MannWhitney U test). **F.** UMAP dimensionality reduction of scRNAseq data from cells isolated from the micro-dissected inner ear region of embryos after Crispr-Cas9 knockdown of either *tyrosinase* (control) or *wls*. Upper-left panel: all cells. Upper-right panel: all cells with expression of the OV marker *stm* overlain as a heatmap. Lower-left panel: all cells in the *stm* OV cluster, colored by subcluster. Lower-right panel: all cells in the OV cluster, with expression of the dorsal OV marker *lmx1bb* overlain as a heatmap, highlighting the pillar-forming cell clusters. **G.** Pseudobulk analyses of *notch1b* in cells from embryos upon knockdown of *tyrosinase* (white) or *wls* (grey) in the whole OV or in the dorsal OV clusters. n denotes the number of biological replicates. Statistical significance was determined using DESeq2, and adjusted P values (Benjamini–Hochberg correction) are indicated. **H.** Representative images of 3D-rendered OVs from membrane-mNeonGreen-expressing embryos at 72 hpf after soaking for 24 hours in either DMSO or the notch inhibitor RO4929097, with pillar region outlined in yellow. Right plot: quantification of OV lumen volume (pL) after 24 hrs of soaking in either DMSO (white) or RO4929097 (grey). P-values as labeled (unpaired, two-tailed Student’s t-test). **I.** Representative images of OVs from membrane-mNeonGreen-expression embryos at the developmental stages indicated, upon knockdown of either *tyrosinase* or *wls*. Prior to imaging at 52 hpf, embryos were injected pericardially with α-Bungarotoxin conjugated to Alexa Fluor 647 to visualize permeabilization of the OV epithelium barrier from 52 to 56 hpf. Right upper plot: quantification of OV lumen volume (pL) at 52 and 56 hpf after knockdown of either *tyrosinase* or *wls*. Right lower plot: normalized intensity (arbitrary units, A.U.) of α-Bungarotoxin-Alexa Fluor 647 in OVs upon *wls* knockdown from 52 hpf (blue) to 56 hpf (red), demonstrating increased permeability of the OV epithelial barrier. P-values as labeled (two-tailed paired t-test and unpaired, two-tailed Student’s t-test). **J.** Representative images of the dorsal OV region of Tight Junction Protein 1a, or ZO-1a (Tjp1a)-tdTomato-expressing embryos after treatment by soaking in either DMSO or the porcupine inhibitor C59. Right upper plot: change in mean intensity (A.U.) of Tjp1a-tdTomato at cell vertices after 24 hours of soaking in either DMSO (white) or C59 (grey), quantified at 48 and 72 hpf. Right-hand lower plot: change in mean intensity (A.U.) of Tjp1a-tdTomato along cell edges after 24 hours of soaking in either DMSO (white) or C59 (grey), quantified at 48 and 72 hpf. P-values as labeled (unpaired, two-tailed Student’s t-test). “n” denotes the number of embryos unless otherwise specified. Scale bar: 50 μm. Data are mean±s.d.

Using a previously described CRISPR-generated *wls* mutant^36^, we found that 46.6% of embryos exhibited early patterning defects, while the remaining embryos displayed normal early development but showed reduced otic vesicle size at later stages, despite undergoing bud initiation, extension, and formation of all pillars (Supp Fig. 4F). Consistent with this, CRISPR–Cas9–mediated *wls* knockdown (*wls* CRISPant embryos) primarily affected later stages, with minimal impact on early patterning (Fig. 4D), likely reflecting partial rescue from maternally deposited Wls protein^35^. To further distinguish early versus late roles of Wnt signaling, we used pharmacological inhibition with C59, a Porcupine inhibitor that blocks Wnt ligand palmitoylation and secretion, allowing temporal control of broad pathway inhibition^37^. Treatment from 24–48 hpf disrupted early patterning of the otic vesicle (Supp Fig. 4G), whereas treatment from 48–72 hpf preserved early patterning but specifically impaired later vesicle expansion (Supp Fig. 4H), resulting in smaller vesicles that nevertheless formed buds and pillars.

Because both genetic and pharmacological perturbations indicated a specific late requirement for Wnt signaling in vesicle inflation, we focused our quantitative analyses on *wls* CRISPant embryos that retained normal early otic vesicle patterning. In these embryos, otic vesicle size was comparable to that of controls up to ∼54 hpf (Supp Fig. 4I). However, vesicles subsequently failed to increase in size, resulting in reduced otic vesicle volume at later stages (Supp Fig. 4I). This reduction could in principle reflect either decreased tissue growth or impaired fluid accumulation within the lumen. To distinguish between these possibilities, we used our U-net–based segmentation pipeline (Fig. 1C) to separately quantify lumen and tissue volumes. Tissue volume was only modestly reduced compared with controls, indicating that epithelial growth was largely preserved (Fig. 4E). In contrast, lumen volume was disproportionately decreased, accounting for most of the reduction in total otic vesicle volume (Fig. 4E). Consistent with this, the tissue fraction (tissue volume/total volume) was significantly higher in Wnt-perturbed embryos than in controls, Together, these results show that the reduced otic vesicle size in Wnt-perturbed embryos results primarily from impaired luminal fluid accumulation rather than a general defect in tissue growth, consistent with reduced hydrostatic pressure within the vesicle lumen.

To identify pathways downstream of Wnt signaling that regulate luminal hydrostatic pressure, we performed single-cell RNA sequencing (scRNA-seq) on micro-dissected inner ear regions from control and *wls* CRISPant embryos. We used *wls* CRISPant embryos because they retain largely normal early patterning while displaying robust defects in vesicle inflation, allowing us to focus on molecular changes associated with the pressure phenotype. Following initial clustering of the combined dataset, otic vesicle cells were identified based on expression of the otic epithelial marker *stm*^38^ and subsequently subset for further analysis. The otic vesicle population was then re-clustered, and the non-sensory dorsal epithelial populations that give rise to canal structures were identified by the expression of *lmx1bb*^16^ (Fig. 4F). Pseudo-bulk analyses were then performed on both the dorsal epithelial level and the whole otic epithelial level. These pseudo-bulk analyses revealed a consistent reduction in *notch1b* expression in *wls* CRISPant embryos (Fig. 4G). This reduction was further validated by HCR-FISH in both *wls* CRISPant embryos and *wls* mutants (Supp. Fig. 5.A-B). Previous studies have demonstrated functional crosstalk between Wnt and Notch signaling during early inner ear development in other vertebrate species^39,40^, and we hypothesized that this relationship extends to later stages of otic vesicle development in zebrafish. Consistent with this hypothesis, pharmacological inhibition of Notch signaling from 48–72 hpf phenocopied major aspects of the *wls* CRISPant phenotype, including reduced vesicle inflation (Fig. 4H). In addition, our previous work demonstrated that prolonged Notch inhibition from 24–72 hpf disrupts the patterning of semicircular canal-budding cells in the otic vesicle^16^. Together with the early budding-cell patterning defects shown above and loss of the lateral crista observed following Wnt inhibition (Supp Fig. 4G), these findings support that Notch signaling is an important downstream effector of Wnt signaling during otic vesicle morphogenesis.

**Figure 5.**
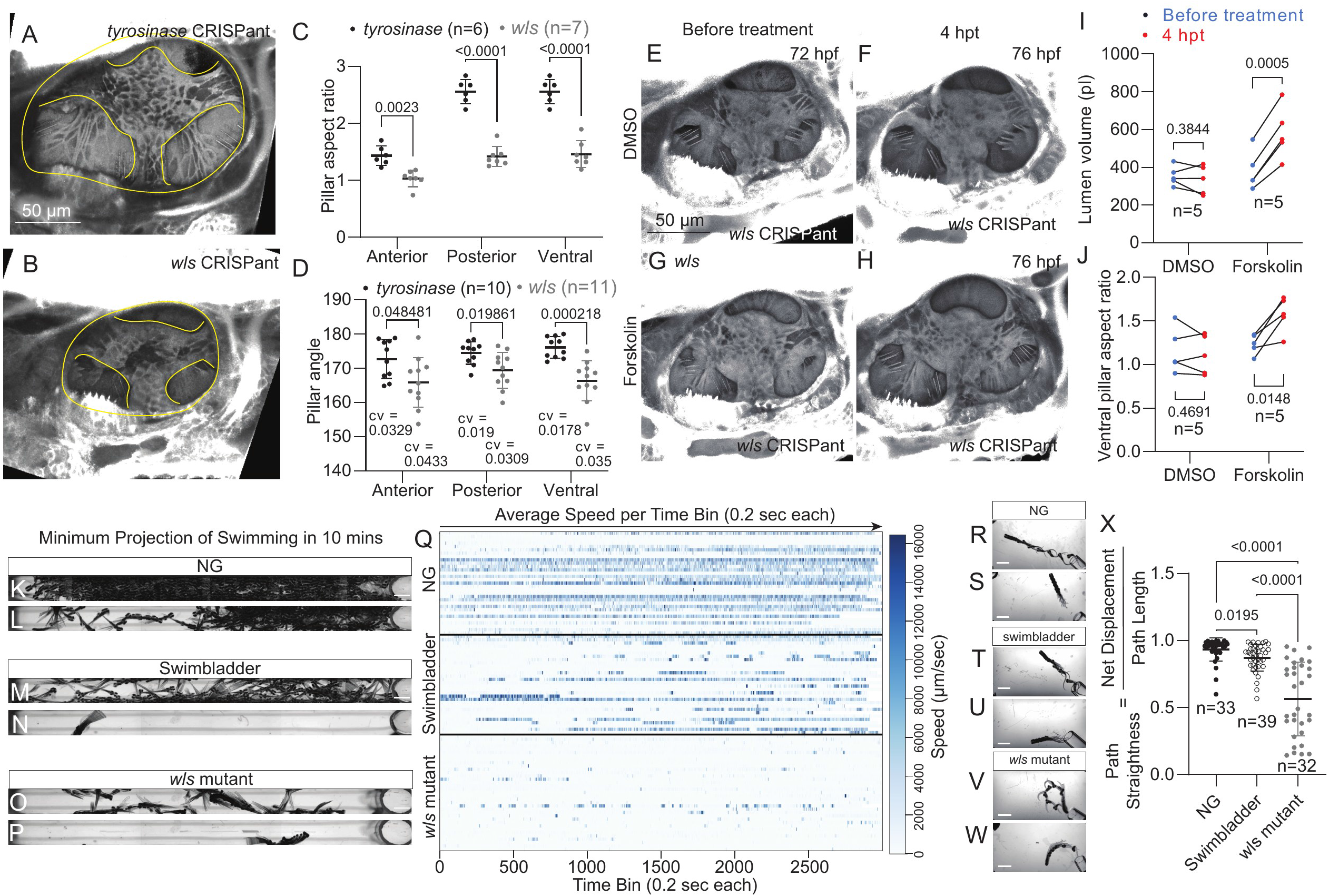
Wnt signaling is required for proper canal form and function. A-B. Representative image of an OV in a *tyrosinase* CRISPant(control) (A) and a *wls* CRISPant (B). **C.** Quantification of pillar aspect ratio of *tyrosinase* (control, black) CRISPants and *wls* (grey) CRISPants by pillar type indicated. **D**. Quantification of pillar angle of *tyrosinase* (control, black) CRISPants and *wls* (grey) CRISPants by pillar type indicated. Coefficient of variation (cv) indicated below each dataset, calculated as the standard deviation from the mean divided by the mean. **E-F.** Representative images of an OV in a *wls* CRISPant before (E) and 4 hpt (F) with DMSO by soaking. **G-H.** Representative images of an OV in a *wls* CRISPant before (G) and 4 hpt (H) with Forskolin by soaking. **I-J.** Quantification of OV lumen volume (pL) in *wls* CRISPants before and 4 hpt with either DMSO (I) or Forskolin (J). P-values as labeled (two-tailed paired t-test). **K-L.** Two representative minimum projections of membrane-mNeonGreen (NG control) 5 dpf larvae swimming in the MCAM channel for 10 minutes. **M-N.** Two representative minimum projections of uninflated swimbladder membrane-mNeonGreen (swimbladder control) 5 dpf larvae swimming in the MCAM channel for 10 minutes. **O-P.** Two representative minimum projections of *wls*^-/-^; membrane-mNeonGreen (*wls* mutant) 5 dpf larvae swimming in the MCAM channel for 10 minutes. Q. Kymograph of average swimming speed in 0.2-second time bins over time for every NG control, swimbladder control, and *wls* mutant larva assayed. Each row represents one larva. **R-S.** Two representative minimum projections of membrane-mNeonGreen (NG control) 6 dpf larvae swimming in the startle response chamber for 308 milliseconds in response to a backwards water current startle trigger. **T-U.** Two representative minimum projections of uninflated swimbladder membrane-mNeonGreen (swimbladder control) 6 dpf larvae swimming in the startle response chamber for 308 milliseconds in response to a backwards water current startle trigger. **V-W**. Two representative minimum projections of *wls*^-/-^; membrane-mNeonGreen (*wls* mutant) 6 dpf larvae swimming in the startle response chamber for 308 milliseconds in response to a backwards water current startle trigger. **X.** Quantification of swimming path straightness in response to a backwards water current startle trigger in NG controls, swimbladder controls, or *wls* mutants. Path straightness defined as the net displacement in 308 milliseconds divided by the total path length traversed over 308 milliseconds. P-values as labeled (Kruskal-Wallis test). “n” denotes the number of embryos. Scale bar: 50 μm. Data are mean±s.d. P-values as labeled (unpaired, two-tailed Student’s t-test, unless otherwise specified). Scale bars in K-P and R-W are all 2 mm.

To gain further insight into the mechanism underlying the reduced vesicle inflation phenotype, we next examined genes involved in fluid accumulation within the otic vesicle lumen. Given the established role of ion transporters in generating osmotic gradients that drive fluid influx^13^, we examined ion transport–related gene expression at both the whole otic vesicle and dorsal epithelium levels. Surprisingly, our pseudo-bulk differential expression analyses revealed that multiple ion transporters known to be involved in zebrafish inner ear development^41^, including *slc12a2* and *kcnq1.1*, were upregulated in *wls* CRISPant embryos (Supp Fig. 5C). This upregulation was further validated using HCR-FISH in both *wls* CRISPant embryos and *wls* mutants (Supp Fig. 5D-G), confirming increased expression at the tissue level. These findings argue against a model in which reduced vesicle pressure results from impaired ion-transport–mediated fluid influx. Instead, they suggest that alternative mechanisms must underlie the loss of hydrostatic pressure. Because vesicle inflation depends not only on ion-driven fluid influx but also on the retention of ions within the lumen, it requires an intact epithelial barrier^42,43^. Given the established roles of Wnt signaling in maintaining epithelial organization^44,45^ and barrier integrity^46^, we hypothesized that Wnt signaling preserves the permeability barrier of the otic epithelium. Loss of barrier function would permit ion leakage, dissipating the osmotic gradient required to drive and maintain luminal fluid accumulation^29^. We tested this directly using two independent approaches: assessing barrier integrity by dye exclusion, and evaluating tight junction organization via ZO-1 localization^20^. In control embryos, a fluorescently labeled dye injected into the periotic space was excluded from the lumen, indicating an intact epithelial barrier (Fig. 4I). In contrast, Wnt-perturbed embryos exhibited leakage of dye into the otic vesicle lumen, indicating compromised epithelial integrity (Fig. 4I). Consistent with this, the tight junction protein ZO-1 localized continuously along the apical junctions of the otic epithelium, but exhibited lower signal in the C59-treated condition (Fig. 4J), consistent with compromised epithelial barrier function. Notably, our scRNAseq analysis did not reveal a significant reduction in the transcription of junctional components in *wls* CRISPant embryos, suggesting that epithelial barrier defects are not primarily driven at the transcriptional level. Instead, the observed decrease in barrier integrity is likely mediated through protein-level regulation of ZO-1 and potentially other junctional components. These results suggest that Wnt signaling is required to maintain epithelial integrity, thereby enabling fluid retention and pressure buildup within the vesicle.

### Wnt signaling is required for proper canal form and function

We next sought to examine whether inhibition of Wnt signaling, by reducing hydrostatic pressure and deflating the otic vesicle, affects post-fusion pillar morphology and canal function. To determine how reduced vesicle expansion impacts canal morphology, we quantified pillar geometry at 72 hpf, when semicircular canal architecture is largely established. In Wnt-perturbed embryos, pillars exhibited significantly reduced aspect ratios (length/width), remaining shorter and wider compared to controls (Fig. 5A-C). In addition, newly formed pillars failed to refine their geometry over time and exhibited greater variability in orientation, as quantified by a larger coefficient of variation in pillar angles (Fig. 5D). These defects indicate that reduced vesicle expansion in Wnt-perturbed embryos results in disrupted pillar geometry and a failure to resolve geometric variability.

To test whether restoring fluid accumulation could rescue these defects, we pharmacologically increased vesicle inflation using forskolin^22^. Forskolin treatment partially rescued pillar morphology in Wnt-perturbed embryos, leading to increased pillar aspect ratios (Fig. 5E-J). Although not fully restored to control levels, this partial rescue supports the idea that impaired vesicle expansion, and consequently reduced hydrostatic pressure, is a primary driver of the observed morphological defects.

Because *wls* mutants exhibit reduced otic vesicle size and abnormal pillar morphology, we asked whether these structural defects are accompanied by vestibular dysfunction, a well-established consequence of disrupted canal formation in zebrafish^47,48^. We assayed two complementary behaviors at 5-6 dpf, an age at which both readouts are reliable: spontaneous swimming behavior, which reports on general locomotor activity^49^, and the startle response, a stereotyped escape reflex that is more directly diagnostic of vestibular function^50–52^. To restrict our analysis to defects arising from later morphogenetic stages rather than earlier patterning abnormalities (Supp Fig. 4F), we selected *wls* mutant larvae that lacked early otic patterning defects for behavioral assays. Spontaneous swimming was assayed using a Multi-Camera Array Microscope (MCAM)^53^ (Supp Fig. 6L), which allowed individual larvae to swim freely within long channels while their trajectories were tracked over extended recordings (Fig. 5K-P) (Supp. Video 2). We used membrane-mNeonGreen (NG) larvae as a baseline control, as this is the genetic background on which the *wls* mutant line used for behavioral assays was maintained. Because *wls* mutants fail to inflate their swim bladders^35^, we could not exclude the possibility that any locomotor defect simply reflected impaired buoyancy rather than vestibular dysfunction. To control for this, we raised a cohort of sibling larvae in submerged holding chambers that physically prevented swim bladder inflation, generating swimbladder-uninflated controls that were otherwise wild-type for inner ear development^54–58^. Brightfield imaging confirmed that NG control, swim bladder-uninflated, and *wls* mutant larvae were grossly similar in overall morphology, with the most consistent differences being the absence of swim bladder inflation and, in *wls* mutants, occasional reduced body length or variable body curvature, along with reduced otic vesicle size and altered canal architecture (Supp. Fig. 6C–K). Kymographs of swimming speed over time for individual fish revealed that *wls* mutants spent the majority of each recording nearly motionless, in contrast to the sustained, high-speed swimming bouts seen in NG controls (Fig. 5K-Q, Supp. video 2). Swim bladder-uninflated controls displayed an intermediate phenotype, with reduced swimming activity relative to NG controls but substantially more sustained movement than *wls* mutants (Fig. 5K-Q, Supp. video 2). Consistently, the cumulative distribution of average swimming speed showed that *wls* mutants were most enriched in the lower-speed bins, followed by swim bladder–uninflated controls and then NG controls (Supp Fig. 6M,N). Together, these results indicate that loss of buoyancy alone partially reduces swimming activity but does not account for the severity of the locomotor defect in *wls* mutant larvae.

To assess vestibular function, independent of general locomotor activity, we next turned to the startle response a rapid, stereotyped escape behavior to visual, audio, or sensory stimuli^50–52^, which we elicited by a directional water current delivered via pipette in a customized container (Supp Fig. 6O-P). NG and swimbladder-uninflated control larvae reliably responded with a fast C-shaped tail-flick that propelled them along a largely straight trajectory away from the source of flow (Fig. 5R-U, Supp. Video 3), visualized by time-colored tracks of individual escape responses (Supp Fig. 6Q-R). In contrast, while *wls* mutants retained the ability to generate a rapid startle reflex, indicating that the underlying sensory cells and motor circuitry remain functional, their escape trajectories frequently looped or curved back on themselves rather than tracing a straight path away from the stimulus (Fig. 5V-W, Supp Fig. 6S, Supp. Video 3). We calculated path straightness for each fish as the ratio of net displacement to total path length, providing a quantitative measure of circling versus directed swimming. Path straightness was modestly but significantly reduced in swimbladder-uninflated controls relative to NG controls, indicating a minor contribution of buoyancy to swimming trajectory (Fig. 5X). *wls* mutants, however, showed a much larger reduction in path straightness relative to both controls, reflecting frequent circling trajectories rather than the persistent, directed swimming seen in controls (Fig. 5X) and supporting a vestibular deficit rather than a nonspecific locomotor defect. Together, these results indicate that Wnt-dependent defects in pillar and canal geometry are accompanied by functional impairment of the vestibular system.

## Discussion

Hydrostatic pressure has long been recognized as a central regulator of tissue morphogenesis in plants, where turgor pressure drives cell and organ growth^59,60^, and has more recently emerged as an important physical regulator in animal developmental systems as well^28,43^, with prior work establishing its role in driving lumen expansion, including in the early otic vesicle formation in zebrafish^13^ and blastocyst formation in mice^29^. In lumenized tissues, hydrostatic pressure is generated as ion transport drives water influx into a confined epithelial lumen, causing tissue inflation until the resulting pressure balances further fluid entry^28^. Here, we extend these concepts by showing that this pressure not only contributes to lumen expansion, but also plays a critical role in regulating epithelial architecture during semicircular canal morphogenesis. Specifically, acute perturbation of vesicle inflation, a proxy for luminal hydrostatic pressure, reciprocally alters pillar aspect ratio, a relationship captured by a physical model of a pressurized epithelial shell that predicts pillar geometry from lumen inflation state. We further show that Wnt signaling sustains luminal hydrostatic pressure by maintaining epithelial barrier integrity. Disrupting Wnt signaling deflates the otic vesicle and reduces pillar aspect ratio, a defect partially rescued by pharmacologically restoring vesicle inflation. Critically, this geometric defect translates into functional consequences: disrupting Wnt signaling also impairs vestibular-dependent swimming behavior, linking canal form to its function. Together, these findings establish hydrostatic pressure as a key mechanical determinant of epithelial pillar geometry, and thus of semicircular canal architecture and vestibular function.

Beyond shaping pillar geometry, we identify hydrostatic pressure as a mechanism for canalizing tissue shape by buffering developmental variability to ensure a reproducible outcome, a feature common among developing tissues^61–63^ that has remained difficult to mechanistically dissect. Newly formed pillars display substantial variability in curvature, reflecting the variability of bud fusion angles. Sustained hydrostatic pressure progressively straightens these pillars, converging initially heterogeneous geometries toward a common, stereotyped morphology. This canalizing role adds to the known morphogenetic functions of hydrostatic pressure, such as controlling tissue size, bending epithelia, and serving as a biochemical and mechanical cue^43,64^, and complements previously described mechanisms of robust development in this system^18^. Because many tissues are sculpted from a pressurized lumen, this pressure-driven error correction may represent a broadly relevant strategy for canalizing organ geometry.

Our physical model provides a mechanistic framework for understanding how ion-driven lumen inflation is coupled to pillar morphogenesis, but several simplifying assumptions warrant consideration. First, the model treats key material properties, including tissue elasticity and viscosity, as fixed parameters, whereas these properties are likely regulated dynamically during development through changes in epithelial organization, cytoskeletal remodeling, and extracellular matrix composition^65–67^. Incorporating feedback between tissue mechanics, fluid transport, and developmental signaling will therefore be an important direction for future models. Second, several parameters, including tissue viscosity, elastic modulus, and luminal pressure, are very demanding to measure *in vivo* and were therefore estimated from published measurements or constrained by fitting experimental dynamics. Future advances in *in vivo* biomechanical measurements will enable more stringent validation and refinement of the model. Despite these limitations, the model successfully reproduces both the acute response to pressure perturbation and long-timescale developmental changes using a single biophysical framework, providing a quantitative foundation for investigating how hydrostatic forces and tissue mechanics interact to shape epithelial architecture.

A key mechanistic insight from our work is the role of Wnt signaling in maintaining epithelial barrier integrity to regulate hydrostatic pressure in the otic vesicle. While the Wnt signaling pathway is well known for its roles in cell fate specification and proliferation^68,69^, its involvement in epithelial permeability and barrier function has been less explored in the context of morphogenesis^46^. Our data support a model in which Wnt signaling preserves epithelial barrier integrity, thereby preventing fluid and ion leakage from the lumen and enabling the buildup of hydrostatic pressure required for normal vesicle inflation. Consistent with previous studies demonstrating functional crosstalk between Wnt and Notch signaling during early inner ear development^39,40^, pharmacological inhibition of Notch signaling phenocopied the reduced vesicle inflation observed following Wnt inhibition, supporting the idea that Notch acts downstream of Wnt during this process. While Wnt inhibition reduced ZO-1 protein abundance, we detected no corresponding transcriptional changes in junctional components, suggesting that Wnt-Notch signaling regulates barrier integrity through post-transcriptional mechanisms or junctional organization rather than transcriptional control of junctional genes. Likewise, the unexpected upregulation of ion transporters following Wnt inhibition argues against impaired ion transport as the primary cause of reduced hydrostatic pressure and instead likely reflects a compensatory response to diminished luminal pressure.

Our findings may also have broader relevance to mammalian inner ear development. In particular, Najarro et al. reported that conditional knock-out of Wls in mouse embryo inner ear results in cochlear duct extension failure^71^. Although it has not been definitively established that hydrostatic pressure drives cochlear duct elongation, this phenotype is consistent with a model in which impaired epithelial barrier function disrupts pressure homeostasis, thereby compromising tissue extension. This observation raises the possibility that pressure-mediated mechanisms identified in the zebrafish otic vesicle may be conserved, at least in part, in mammalian systems.

An additional layer of regulation may involve pressure relief mechanisms within the otic vesicle. The endolymphatic duct and sac have been proposed to function as a pressure valve, modulating luminal hydrostatic pressure to prevent excessive inflation^72^. Whether this structure plays a role during the pillar morphogenesis stages studied here remains untested. In this context, the balance between fluid secretion, epithelial barrier integrity, and pressure release may represent a key regulatory mechanism governing tissue shape. Directly measuring luminal hydrostatic pressure in both wild-type and perturbed conditions and determining whether the endolymphatic duct contributes to its regulation during canal morphogenesis, will be essential to quantitatively test this model and validate the predicted mechanical relationships.

Finally, our findings may have implications for human disease. In our model, *wls* mutants with reduced otic vesicle size exhibit behavioral abnormalities, including circling swimming patterns, suggestive of impaired vestibular function. This observation is consistent with the essential role of the inner ear in maintaining balance and spatial orientation and supports the idea that disrupted morphogenesis can lead to functional deficits. In humans, disorders such as Ménière’s disease are characterized by vertigo, hearing loss, and tinnitus and have long been associated with dysregulation of inner-ear fluid homeostasis and accumulation of endolymphatic fluid (endolymphatic hydrops)^73^. Emerging evidence further suggests that barrier dysfunction and impaired ion/fluid regulation may contribute to disease pathophysiology^74^. In this context, our findings highlight how hydrostatic pressure–driven morphogenesis shapes organ geometry, and how disruptions to these physical processes may ultimately compromise organ function.

In summary, we propose a model in which Wnt-dependent regulation of epithelial barrier function controls luminal hydrostatic pressure, which in turn governs epithelial morphology and ensures robust morphogenesis. This work highlights the importance of integrating biochemical signaling with physical forces to understand developmental processes and suggests that hydrostatic pressure may serve as a general mechanism for coordinating tissue shape and physiological functions across organ systems.

## Materials and methods

### Animals

Zebrafish (*Danio rerio*) of the EK wild-type strain were used in this study. Adult fish were maintained under a 14-h light/10-h dark cycle, and embryos were obtained through mating of adult males and females (3–18 months old) and raised in egg water at 28.5°C. The following transgenic and mutant lines were used: *Tg(actb2:membrane-neongreen-neongreen)*^14^, *TgKI(tjp1a-tdTomato)^pd^*^1224^ ^75^, and a CRISPR-generated *wls* mutant line obtained from Eric Liao’s laboratory carrying a 7-bp deletion in exon 2 of *wls*^36^, as determined by Sanger sequencing. Because homozygous *wls* mutants fail to survive to adulthood, the line was maintained as heterozygous carriers. To visualize the inner ear morphology in *wls* mutants, the *wls* mutant line was crossed with *Tg(actb2:membrane-neongreen-neongreen)* fish. Offspring carrying both the *wls* mutation and the membrane-mNeonGreen (NG) transgene were identified by genotyping and subsequently incrossed to generate membrane-mNG-positive homozygous *wls* mutant embryos for confocal imaging experiments. Homozygous mutant embryos were screened by ear and swim bladder phenotype. Genotyping was performed by heteroduplex mobility shift assay^76^ using the following primers: forward, 5’-ctatctggccaccaagtgtgt-3’; reverse, 5’- cagtgtgtgtgagcttatggct-3’. All animal experiments were conducted in accordance with protocols approved by the institutional regulatory review at Duke University School of Medicine.

### Otic vesicle segmentation

The otic vesicle (OV) segmentation pipeline was developed using the PyTorch framework^77^. Training data were generated through semi-automated three-dimensional annotations using the segmentation tool in ITK-SNAP^78^. A 3D U-Net architecture was implemented using the segmentation_models_pytorch_3d package^79^, with a ResNet-34 backbone employed as the encoder network^80^. To improve adaptability across experimental conditions, the segmentation framework incorporated both continual learning and transfer learning modules. Depending on experimental requirements, the model was trained to perform segmentation using the periotic dye imaging channel. All scripts used for model training and data processing are publicly available on GitHub (https://github.com/MunjalLab/OV_Segmentation).

### Genome editing by CRISPR-Cas9

#### *wls* CRISPant embryo

To generate *wls* CRISPant embryos, single guide (sg) RNA targeting *wls* was co-injected with Cas9 protein at the one-cell stage as previously described^81^. Injection mixtures contained 200 ng/μL sgRNA and 500 ng/μL Cas9 protein (PNA Bio, CP-03). Unless otherwise specified, embryos were screened at 72 hours post-fertilization (hpf) for reduced OV size with normal SCC patterning prior to downstream experiments. The following target sequence was used for sgRNA generation: 5′-GTAGGCCAGCCTGACATCAATGG-3′.

#### *tyrosinase* CRISPant embryo

Control embryos were generated by injection of sgRNA targeting tyrosinase together with 500 ng/μL Cas9 protein at the one-cell stage, as previously described by Sorlien et al^81^. Injection mixtures contained 200 ng/μL sgRNA. *Tyrosinase* CRISPant embryos were screened for loss of pigmentation prior to each experiment and subsequently used for live confocal imaging or hybridization chain reaction fluorescence *in situ* hybridization (HCR-FISH) analysis. The following target sequence was used for sgRNA generation: 5′-GGACTGGAGGACTTCTGGGG-3′.

### Drug treatments

For forskolin treatment, embryos were dechorionated and incubated in egg water containing either 50 μM or 70 μM forskolin. Forskolin (Cayman Chemical, 1101810) was dissolved in DMSO and stored as a 50 mM stock solution at −20 °C. To assess pillar shape changes, embryos were treated with 50 μM forskolin from 54–58 hpf, 60–64 hpf, or 66–70 hpf. To measure pillar angle changes, embryos were treated with 70 μM forskolin for 1 h at 60 hpf. To test whether forskolin treatment rescues the *wls* CRISPant phenotype, *wls* CRISPants were treated with 50 μM forskolin for 4 h at 72 hpf. In all experiments, embryos were imaged before and after treatment. Control embryos were treated in parallel with equivalent concentrations of DMSO in egg water.

Hyaluronidase (HAase) treatment was performed similarly to previous work^14^. Embryos were first dechorionated, anesthetized by soaking in 1X tricaine, screened for bud touching, and then mounted dorsally in a canyon mount cast with 1.5% agarose dissolved in egg water for injection. Embryos were then immobilized by cardiac injection with 500 uM AlexaFluor647-alpha-bungarotoxin (aBt) (stock: *Bungarus multi cinctus* aBt from ThermoFisher cat. no. B35450, 1 M). Embryos were then screened for newly-formed pillars, evident by formation of a passage between apposing buds, and imaged at least 1 hour after aBt injection. Embryos were then injected with HAase. HAase injection solution was made from 200 unit/ml HAase diluted in 1x PBS and 0.5% Phenol Red (stock: Streptomyces HAase from Sigma, 3000 unit/ml). Solution was injected into the periotic space (anterior and posterior regions surrounding the OV). Control injection solution was made with 1x PBS and 0.5% Phenol Red without HAase. Embryos were imaged again after injection with HAase, roughly 1 hour after the first image.

For Wnt inhibition, embryos were dechorionated and incubated in egg water containing 1 μM C59 from 24–48 hpf or from 48–72 hpf. C59 (Sigma, 5004960001) was dissolved in DMSO and stored as a 1 mM stock solution at −80 °C. Control embryos were treated in parallel with 1% DMSO in egg water. Embryos treated from 24–48 hpf were imaged at both 48 hpf and 72 hpf. After treatment, embryos were washed thoroughly with egg water and transferred to fresh egg water prior to continued incubation. Embryos treated from 48–72 hpf were imaged at 72 hpf. To quantify changes in ZO-1 intensity following C59 treatment, ZO-1 reporter embryos (*TgKI(tjp1a-tdTomato)^pd12^*^24^) were first imaged at 48 hpf prior to treatment, then individually transferred to 48-well plates (one embryo per well) containing 1 μM C59. The same embryos were re-imaged at 72 hpf following treatment.

For Notch inhibition, embryos were dechorionated and incubated in egg water containing 1 μM RO4929097 from 48–72 hpf, similar to how previously described^16^. RO4929097 (ApexBio, A4005) was dissolved in DMSO and stored as a 50 mM stock solution at −20 °C. Control embryos were treated in parallel with 0.1% DMSO in egg water. Embryos were imaged at 72 hpf following treatment.

#### Two-photon ablation

Ablation of cells within the OV was performed using a Mai Tai HP two-photon laser system (Spectra-Physics, Santa Clara, CA) similar as previously described^14^. The laser was tuned to 800 nm and operated at 50% power. The confocal pinhole was fully opened, and targeted cells were ablated using repeated spot scanning for 200–1,000 iterations. Successful ablation resulted in disruption of the otic epithelium and subsequent collapse of the OV. To examine the morphological change following the ablation, the embryos were imaged within 30 min post-ablation at various developmental stages. To capture the fluid dynamics after ablation, the ablated embryos were imaged every 3 min post-ablation for 30 min.

#### Permeability assay

To assess OV permeability, 500 mM α-Bungarotoxin-AF647 (Invitrogen, B35450) was injected into the pericardial cavity of *wls/tyrosinase* CRISPants at 52 hpf. Following injection, embryos were immediately imaged to acquire baseline OV fluorescence intensity at 52 hpf. The same embryos were re-imaged at 56 hpf to evaluate changes in OV permeability during lumen collapse in *wls* CRISPants. For each experimental replicate, the same injection needle was used for both tyrosinase control embryos and *wls* CRISPants to minimize variability in injection volume. The OV lumen was segmented using the U-Net–based image segmentation pipeline, and the lumen mean fluorescence intensity and lumen volume were quantified for each embryo based on the segmentation results. For normalization, the lumen intensity of each *wls* CRISPant was divided by the average lumen intensity of tyrosinase control embryos from the same experimental replicate at the corresponding time point (0 h post-injection at 52 hpf and 4 h post-injection at 56 hpf).

#### HCR-FISH Assay

Multiplex Hybridization Chain Reaction-fluorescence in situ hybridization (HCR-FISH) was carried out following previously established protocols^14^. Gene-specific HCR probe sets were purchased from Integrated DNA Technologies (IDT; oPools), while hybridization, amplification, and wash buffers together with fluorescent hairpin amplifiers were obtained from Molecular Instruments. Embryos at the desired developmental stages were manually dechorionated and fixed in 4% paraformaldehyde (PFA) prepared in PBS. Following fixation, samples were washed in PBS for 3 times and permeabilized using pre-chilled acetone and equilibrated in hybridization buffer for 30 min at 37°C. Probe hybridization was performed overnight at 37°C in hybridization buffer containing HCR probe sets (1 pmol per probe set). After excess probes were removed through a series of washes, embryos were incubated overnight at room temperature in amplification buffer containing snap-cooled fluorescent hairpins to enable HCR-mediated signal amplification. Samples were protected from light during the amplification step to minimize photobleaching of the fluorescent hairpins. Following amplification, embryos were washed four times with 5× SSC supplemented with 0.1% Tween-20 (SSCT). Samples were subsequently mounted for confocal imaging or maintained at 4°C prior to imaging. For all comparative experiments, control and experimental embryos were processed together under identical conditions, including shared incubation times, reagent concentrations, and handling procedures, in order to minimize technical variability. During image acquisition, confocal settings including laser intensity, detector gain, and exposure parameters were kept constant and below saturation thresholds across samples to permit reliable comparison of fluorescence signal intensity.

#### Immunohistochemistry

For staining with hyaluronan binding protein (HABP), embryos at various stages were dechorionated and fixed with 4% paraformaldehyde (PFA) in PBS, as previously described^14^. To distinguish embryos at distinct stages stained in the same tube, tails were cut with a surgical knife at varying lengths corresponding to different developmental stages. Embryos were permeabilized with pre-chilled acetone as previously described. Blocking was performed with 5% bovine serum albumin (BSA; stock: Sigma-Aldrich cat. no. A3311) in 1x PBS with 0.1% Tween-20 (PBT) at room temperature for 1 hour, rocking and then incubated with primary antibody solution containing biotinylated-HABP (Sigma-Aldrich, 385911; 1:50) in 5% BSA with 1x PBT at 4°C overnight. Embryos were then washed three times with 1xPBS at room temperature and incubated in secondary antibody solution containing streptavidin-AlexaFluor568 (ThermoFisher Scientific, S11226, 1:500), phalloidin-AlexaFluor647 (ThermoFisher Scientific, A22287, 1:200), and Hoescht (Thermo Fisher Scientific, 62249, 1:1000) at 4°C overnight on a rocker, covered. Embryos were then washed three times with 1xPBS rocking at room temperature and then kept in the dark at 4°C until ready to image.

#### Confocal imaging

Confocal images were performed with a Zeiss LSM 980 confocal microscope using a C-Apochromat 40×1.2 NA objective for all fluorescence data as previously described^16^. For live imaging, dechorionated embryos were anesthetized in 1× tricaine and then mounted dorsolaterally using a canyon mount cast in 1.5% agarose dissolved in egg water, submerged in egg water, and covered with a coverslip, as previously described^72^. For fixed imaging, embryos were mounted the same way but submerged in 1xPBS.

#### Timelapse confocal imaging

For timelapse confocal imaging of pillar formation and straightening, embryos were dechorionated, anesthetized by soaking in 1X tricaine, and then mounted dorsally in a canon mount cast with 1.5% agarose dissolved in egg water for injection. Embryos were immobilized by cardiac injection with 500 uM AlexaFluor647-alpha-bungarotoxin (aBt, stock: 1 mM *Bungarus multicinctus* aBt from ThermoFisher cat. no. B35450)^82^. Embryos were mounted for imaging at least 30 minutes after aBt injection in a canon mount cast with 1.5% agarose dissolved in egg water and covered with a no. 1 coverslip (25×25 mm), as previously described^72^. The embryo mounting chamber was kept filled with egg water to minimize impacts of water loss on sample drift. Embryos were filmed with the confocal settings described above in a custom built, temperature-controlled chamber set to 28.5°C, and stacks with a step size of 0.5 µm were acquired every 30 minutes at various developmental stages for up to 20 hours, as previously described^14,18^. Images were processed in Zen software (v3.5) and Fiji software^83^ and 3D rendered in FluoRender^84^ for display. The supplemental video was further annotated in Adobe Premiere Pro.

### Single-cell RNAseq

#### Library preparation

Library preparation was performed as previously described^31^. To minimize cell adhesion to plastic surfaces, 1.5 mL collection tubes were coated with 10% BSA for 10 minutes, rinsed with PBS, and maintained on ice prior to use. Dissections were performed in modified DMEM supplemented with tricaine, consisting of 1× DMEM (11039-021, Life Technologies), 10 mM HEPES (cat), 1× non-essential amino acids (11140076, Life Technologies), 1 mM sodium pyruvate (11360070, Life Technologies), 100 μM β-mercaptoethanol (M3148, Sigma), and 0.33 mg/mL tricaine (E10521, Sigma). For each biological replicate, approximately 30 *tyrosinase* CRISPants and 30 *wls* CRISPants at 72 hpf were collected. Dissections were performed by two researchers using 22-gauge trabecular needles and scalpels. After careful removal of the brain tissue, the inner ear region was isolated and transferred into BSA-coated collection tubes. Following dissection, excess DMEM was removed and 100 μL trypsin-EDTA (25300054, ThermoFisher) was added. Samples were briefly centrifuged, the supernatant was removed, and another 100 μL trypsin-EDTA was added. Samples were incubated at 28°C for 10 minutes, mechanically dissociated by gentle pipetting, and then subjected to two additional rounds of incubation at 28°C for 5 minutes followed by pipetting to further dissociate the tissue into single cells. Subsequently, 100 μL FACSMax Cell Dissociation Solution (AMS.T200100, Amsbio) was added, and samples were gently pipetted up and down while avoiding bubble formation, followed by incubation on ice for 2 minutes. During incubation, fresh BSA-coated tubes were rinsed with PBS. An additional 600 μL FACSMax solution was then added, and the cell suspension was filtered through a 40 μm cell strainer (431750, Corning) into the prepared BSA-coated tubes. Cells were centrifuged at 300 × g at 4°C for 3 minutes in a pre-cooled swinging bucket centrifuge. The supernatant was removed, and cells were resuspended in 1 mL chilled PBS containing 0.5% BSA. Samples were centrifuged again at 300 × g for 3 minutes, after which the supernatant was removed and cells were resuspended in 200 μL pre-chilled 1× PBS. Cell numbers were quantified using a hemocytometer. Cells were then fixed using the Parse Biosciences Evercode WT v3 Low Input Fixation Kit (ECFC3300) and stored at −80°C in a Styrofoam container. In total, four biological replicates were prepared. Fixed cells were subsequently processed for barcoding and library preparation according to the Parse Biosciences Evercode WT v3 protocol (ECWT3300).

### Single cell library sequencing

A total of eight libraries were sequenced by Innomics Inc. (San Jose, CA; RRID:SCR_024849) using the DNBSeq (formerly MGISEQ-2000) platform^85,86^ with paired-end 100 bp (PE100) reads. Sequencing was performed across eight G400 lanes to generate sufficient coverage for downstream single-cell transcriptomic analysis.

### Single-cell RNAseq analysis

To analyze the embryo-wide Daniocell atlas^30,87^, single-cell RNA-seq data corresponding to otic tissue from 40 to 72 hpf were extracted and analyzed using the Seurat package in R^88–90^. Differential gene expression analyses were performed within Seurat, and volcano plots were generated using the EnhancedVolcano package.

For the inner ear–specific single-cell RNA-seq dataset^31^, downstream analyses were performed in Python using the Scanpy package^91^. Standard workflows including data preprocessing, dimensionality reduction, clustering, visualization, and differential gene expression analysis were carried out in Scanpy. Volcano plots were generated using the EnhancedVolcano package in R.

To analyze the *wls* CRISPants single-cell RNA-seq data, raw FASTQ files were processed and aligned against the zebrafish reference genome GRCz11 using the Danio_rerio.GRCz11.dna_sm.primary_assembly.fa genome assembly^92^ and Danio_rerio.GRCz11.111.gtf gene annotation files obtained from Ensembl release 111^93^. Following alignment and generation of gene expression count matrices, downstream single-cell RNA-seq analyses were primarily performed in Python using the Scanpy package^91^. Standard preprocessing steps included quality control, normalization, dimensionality reduction, clustering, and differential gene expression analysis. For pseudobulk differential expression analysis, single-cell counts were aggregated by biological replicate and analyzed in R using the DESeq2 package^94^.

All analysis scripts used in the single-cell RNAseq analyses can be found on GitHub (https://github.com/MunjalLab/scRNAseq_WntProject). Dataset is available on gEAR Dataset Explorer (https://umgear.org/dataset_explorer.html) by searching ‘hydrostatic pressure’.

### Behavioral assays

#### Embryo collection and phenotypic screening

Embryos were obtained from heterozygous *wls* mutant incrosses and maintained in egg water, which was replaced regularly. To generate swim bladder-uninflated controls, a subset of membrane-mNG larvae was transferred at 3 days post-fertilization (dpf) into a submerged 3D-printed holding chamber that prevented access to the air–water interface while allowing water and gas exchange through a mesh-covered lid (Supp Fig. 5A-B), a method previously used to prevent swim bladder inflation in larvae by blocking surfacing behavior^54–58^. Homozygous *wls* mutants were identified at 3–4 dpf based on the characteristic combination of an uninflated swim bladder, reduced OV size, and normally patterned semicircular canal pillars.

#### Multi-Camera Array Microscope (MCAM) assay

Spontaneous swimming behavior was recorded at 5 dpf using a Multi-Camera Array Microscope (MCAM)^53,95,96^. Individual larvae were placed into 4-mm-wide channels of a custom channel well plate. Prior to imaging, larvae were transferred from a 27°C dark incubator to a holding dish at room temperature under ambient light for 20 min, followed by an additional 5 min acclimation in the imaging chamber. Larvae were then recorded for 10 min under brightfield illumination at 10 frames s⁻¹. Mutant and control larvae were imaged in alternating order to minimize potential time-of-day effects. Videos from five adjacent cameras spanning each channel were saved as uncompressed.nc files and converted to MP4 format for downstream analysis. Each day this assay was performed, at least two different groups (from NG, Swimbladder, and *wls* mutant) were assayed to control for any variability due to assay day.

#### Flow-evoked startle assay

Flow-evoked startle responses^50–52^ were recorded at 6 dpf using siblings of the larvae analyzed by MCAM^53,95,96^. Individual larvae were placed within a 3D-printed circular enclosure secured to the bottom of an egg water-filled Petri dish lid, which restricted the larva to the imaging field while allowing controlled delivery of the stimulus. The enclosure contained a notch that stabilized a Pasteur pipette while permitting water exchange. Larvae were oriented with the head facing away from the pipette, and a gentle backward water current was generated by withdrawing water through the pipette to evoke a startle response. Larvae that moved before stimulation were repositioned and retested. Videos were acquired on an Olympus MVX10 microscope equipped with a Hamamatsu ORCA-Spark CMOS camera at 64.9 frames s⁻¹ for 650 frames. Each day this assay was performed, at least two different groups (from NG, Swimbladder, and *wls* mutant) were assayed to control for any variability due to assay day.

### Quantification, theoretical model and statistical analysis

#### Quantification

Volume measurements and tissue fraction quantification

Segmentation of the lumen and whole OV was performed using a U-Net–based convolutional neural network segmentation pipeline. The model used for whole OV segmentation was generated by transfer learning from a previously trained U-Net model developed for membrane-based lumen segmentation. To adapt the model for whole OV segmentation, the pretrained network was fine-tuned using confocal image stacks of the periotic dye channel, and the resulting model was applied to segment the entire OV volume from periotic dye images. Voxels within the segmentation masks were converted to volumetric measurements (pL) based on the calibrated voxel dimensions. To quantify extracellular matrix (ECM) volume, 500 mM α-Bungarotoxin-AF647 was injected into the pericardial cavity to label the periotic space. ECM regions were manually segmented in Imaris and converted to volumetric measurements using voxel size calibration. Tissue fraction was calculated using the following formula:

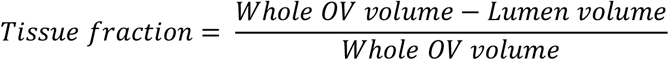

### Pillar aspect ratio measurements

Confocal image stacks of the OV were three-dimensionally rendered using FluoRender^84^. Pillar length and width were measured using the measurement toolkit in FluoRender. Pillar width was defined as the narrowest region located at least one cell diameter away from the fusion zone. Prior to measurement, each pillar was rotated to align parallel to the viewing plane to minimize errors caused by projection artifacts. The pillar aspect ratio was calculated as length divided by width.

### Pillar angle measurements

Confocal image stacks of the OV were opened in the FIJI plugin Mastodon^97^, to enable non-orthogonal reslicing of the 3D data. The images were rotated about the middle of the pillar fusion zone (evident by rounder cell shapes and a dimpling in the pillar epithelium) to find the plane at which the pillar was bisected from one base to the other. Three points were then placed in this position adjusted to the longitudinal axis of the pillar: one at the center of each of the pillar bases, dorsal and lateral (P_D_ and P_L_) and one at the center of the pillar fusion zone (P_F_). The pillar was then rotated to look down the length of the pillar from each base to adjust these base coordinates to the radial axis of the pillar. These 3D coordinates were then saved to a csv file from Mastodon and analyzed in MatLab to calculate pillar angle. Pillar angle was calculated using vectors connecting the 3D coordinates with the following formula:

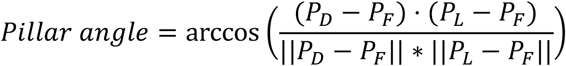

All analysis scripts used for angle calculations can be found on GitHub. (https://github.com/MunjalLab/wls_general_analysis)

### Timelapse data analysis

Frames of timelapse data were cropped to timepoints from pillar fusion through pillar straightening. OV lumen volume and anterior pillar angle were quantified at each timepoint. To account for variability in initial conditions, both lumen volume and pillar angle were normalized with the following methods. Lumen volume was normalized by dividing values by the initial volume value, and pillar angle was normalized by dividing values by the potential for straightening (the initial angle subtracted from 180°). Change was computed as the difference in normalized volume or angle value from the first timepoint to each subsequent timepoint. These normalized values were then pooled from all 5 timelapses for statistical analyses. This analysis was also extended to values from the forskolin and ablation experiments to pool these with the wildtype timelapses for statistical analyses.

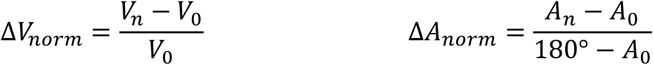

All analysis scripts used for timelapse analyses and plotting can be found on GitHub. (https://github.com/MunjalLab/wls_general_analysis)

### Intensity analysis of multiplex in situ HCR

For HCR-FISH analysis of *slc12a2* and *kcnq1.1*, maximum-intensity projections spanning the entire OV were generated from confocal z-stacks in Fiji software. Regions of interest (ROIs) corresponding to the semicircular canal cristae (SCC), the primary domains expressing these ion transporters, were manually selected. For each embryo, three circular background ROIs were selected within the periotic mesenchyme by ‘Oval selection’ in FIJI. Each background ROI had a diameter of approximately 10 μm, and the ROI size was kept consistent across all embryos within each experiment. Mean fluorescence intensity within the SCC ROI was normalized to the average mean background intensity.

For *notch1b* intensity analysis, maximum-intensity projections were similarly generated for ROI selection. ROIs encompassed the entire OV while excluding neuromasts when present within the projection. For each embryo, three circular background ROIs were selected within the periotic mesenchyme by ‘Oval selection’ in FIJI. Each background ROI had a diameter of approximately 10 μm, and the ROI size was kept consistent across all embryos within each experiment. The mean OV intensity was normalized to the average background intensity.

### Intensity analysis of ZO-1 reporter

To quantify ZO-1 reporter intensity, sum-intensity projections were generated using an identical number of z-slices for all embryos within the same developmental stage in Fiji software. For each embryo, 10 cell edges and 10 cell vertices were randomly selected before and after treatment. Cell edge ROIs were manually generated in Fiji by ‘Segmented line’ with ‘Line width’ set to 2 (0.414 μm), and cell vertices ROIs were manually generated in Fiji by ‘Oval selection’. Mean fluorescence intensities for edges and vertices were calculated separately for each image. Changes in intensity before and after treatment were used for statistical analysis and data visualization.

### Behavioral video processing, tracking, and quantification

MCAM videos were stitched using a custom Python pipeline that registered the five adjacent camera views using a calibration image, generating a continuous view of each swimming channel. The stitched videos were subsequently processed in Fiji software by cropping and rotating the channels to remove boundary artifacts while preserving the available swimming area. Olympus startle-response videos were spatially cropped to remove the enclosure and temporally aligned by manually identifying the onset of the startle response. A 21-frame interval beginning at the first frame of the response was retained for subsequent analysis.

Larval trajectories from both MCAM and Olympus recordings were extracted using FastTrack^98^. Tracking parameters, including image threshold, object size limits, and morphological dilation, were optimized individually for each recording to maximize continuous detection of the larval body while minimizing background noise. All tracking results were visually inspected, and discontinuous trajectories or incorrectly detected objects were manually corrected when necessary. Frame-by-frame head position, body orientation, and additional tracking parameters were exported for downstream analysis.

For MCAM recordings, swimming speed was calculated from the Euclidean displacement between consecutive tracked positions and converted to physical units using the calibrated pixel size and imaging frame rate. Instantaneous speeds were additionally averaged over non-overlapping windows of 3 frames to examine swimming behavior at a higher temporal scale. Swimming behavior was visualized using cumulative distribution functions, speed distributions, and kymographs generated with custom Python scripts.

For the flow-evoked startle assay, only the head coordinates exported from FastTrack^98^ were used for trajectory analysis. Path straightness was calculated as the ratio of net displacement to the total path length traveled during the analyzed response interval, where values approaching 1 indicate directed movement and lower values indicate increasingly curved swimming trajectories. Custom Python scripts were used to calculate path straightness from the exported head coordinates and to visualize larval trajectories from the flow-evoked startle assay.All analysis scripts used in the behavioral analyses can be found on GitHub (https://github.com/MunjalLab/wls_Behavioral-assays).

## Theoretical model

### 1. MODEL DESCRIPTION

#### Geometrical assumptions

We model the morphodynamics of the otic vesicle at an early developmental stage after the formation of a single pillar, corresponding to the fused budding structure. The otic vesicle is assumed to have a spherical geometry with radius *R*_2_, surrounded by a periotic space containing mesenchymal cells. The vesicle is modeled as an epithelial shell of uniform thickness ℎ enclosing a lumen. Within the lumen, we model the pillar as an axisymmetric cylindrical structure of radius *R*_l_, capped at both ends by spherical caps of height *d* (as illustrated in the schematic, Fig. 2K). The interior of the pillar contains a hyaluronan-rich gel network, which is confined by an epithelial wall that is assumed to have the same thickness ℎ as the epithelial shell surrounding the lumen. The entire system is assumed to be axisymmetric, and for simplicity, we consider only a single pillar within the otic vesicle. We assume that throughout the simulated dynamics the otic vesicle remains approximately spherical and the pillar retains the idealized axisymmetric geometry shown in Fig. 2K. Changes in compartment volumes are therefore represented through changes in the corresponding geometric parameters. Under these geometrical assumptions, the volumes of the different compartments are given by:

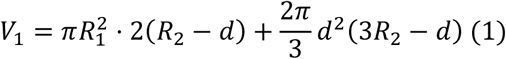

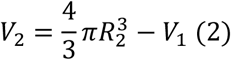

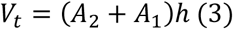

Here, *V*_l_ denotes the volume of the pillar, *V*_2_ is the volume of the otic vesicle lumen excluding the pillar, and *V*_t_is the total epithelial tissue volume. The height of each spherical cap is given by 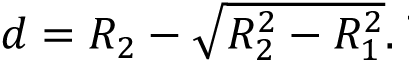. The relevant interfacial areas are defined as follows: The pillar–lumen interface area: *A*_l_ = 2*π R*_l_ ⋅ 2(*R*_2_ − *d*), The lumen–external tissue interface area: *A_2_* =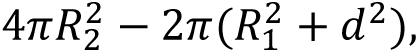, The pillar–external tissue interface area: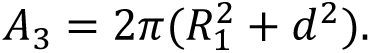.

### Volume dynamics

We assume that all geometrical evolution is driven by fluid transport across the interfaces separating different compartments. Accordingly, the volume dynamics of each compartment are governed by the net water flux across its boundaries:

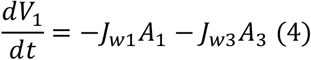

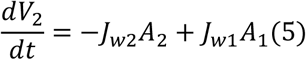

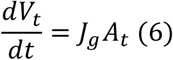

Here, *V*_l_, *V*_2_, and *V*_t_ denote the volumes of the pillar, lumen, and epithelial tissue, respectively. *J*_wl_, *J*_w2_ and *J*_w3_ represent water fluxes across the pillar–lumen interface, lumen–external tissue interface, and pillar–external tissue interface, respectively. We define efflux (flow out of a compartment) as positive. *J*_g_ denotes the effective growth rate of the epithelial tissue, accounting for both lumen wall and pillar wall expansion. *A*_t_ is the total tissue surface area contributing to growth. In the simulations presented here, epithelial tissue growth is neglected (*J*_g_ = 0); thus, Eq. (6) is retained only for completeness. Water transport across each interface obeys the linear hydraulic permeability law that accounts for both hydrostatic and osmotic pressure differences [1]. Specifically, *J*_wl_ = *α*_l_[(*P*_l_ − *P*_2_) − (Π_l_ −Π_2_)], *J*_w2_ = *α*_2_[(*P*_2_ − *P*_O_) − (Π_2_ − Π_O_)], *J*_w3_ = *α*_3_[(*P*_l_ − *P*_O_) − (Π_l_ − Π_O_)]. Here, *P*_i_ and Π_i_ denote the hydraulic and osmotic pressures, respectively, with *i* = 0,1,2 corresponding to the external mesenchymal tissue, pillar, and lumen. The coefficients *α*_i_ represent the hydraulic permeability of the corresponding interfaces. The time evolution of the geometrical variables *R*_l_, *R*_2_ and ℎ is determined implicitly through the volume relations given in Eqs. (1)–(3), which couple fluid transport to shape changes.

### Ion dynamics

We denote the ion concentration in each compartment by *c*_i_, with *i* = 0,1,2 corresponding to the external mesenchymal tissue, pillar, and lumen, respectively. Ion transport in the lumen is governed by fluxes across the surrounding interfaces. The total number of ions in the lumen evolves according to:

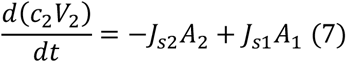

Here, *J*_sl_ and *J*_s2_ denote the ion fluxes across the pillar–lumen interface and the lumen–external tissue interface, respectively. The first term represents ion efflux from the lumen to the external tissue, while the second term accounts for ion exchange between the pillar and lumen. The pillar interior consists of a hydrogel composed of a polymer network (hyaluronan), interstitial fluid, and mobile ions. To describe ion partitioning within the gel, we adopt a standard triphasic theory [2] for charged hydrogels, which yields the following approximation for the ion concentration inside the pillar:

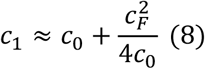

Here, *c*_F_ denotes the fixed charge density associated with the gel network. We assume monovalent ions (valence ±1). This relation corresponds to Donnan equilibrium between the pillar and the external medium, giving rise to an osmotic pressure difference that drives water influx into the gel. Ion transport across interfaces is modeled as a combination of passive diffusion and active transport:

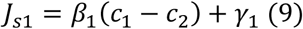

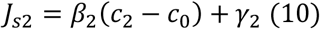

Here, *β*_i_ represents the effective permeability (passive transport coefficient) of ions across each interface, and *γ*_i_ denotes the active pumping rate.

The osmotic pressure in each compartment is computed from the concentration of mobile ions using the van’t Hoff relation.

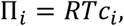

where *R* is the universal gas constant, *T* is the absolute temperature, and *c*_i_ is the concentration of mobile ions in compartment *i*. In the pillar, the elevated ion concentration resulting from Donnan partitioning (Eq. 8) increases the osmotic pressure, promoting water influx into the HA-rich hydrogel.

### Mechanics

To determine the hydraulic pressures in each compartment, we incorporate the mechanical response of the surrounding tissues. We model both the lumen wall and pillar wall as viscoelastic materials described by a Maxwell constitutive relation. The evolution of the circumferential surface stress in each structure is governed by:

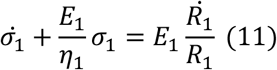

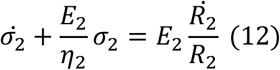

Here, *σ*_l_ and *σ*_2_ denote the circumferential surface stresses in the pillar wall and lumen wall, respectively. *E*_i_ and *η*_i_ are the effective elastic modulus and viscosity of the tissue, and *R*_i_ are the corresponding radii. For both structures, we define a reference radius *R*_i,O_ corresponding to a stress-free configuration. The hydraulic pressure difference across the lumen wall is determined by the balance of surface tension and curvature (Laplace law):

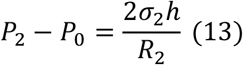

The pillar interior is modeled as a hydrogel composed of a solid polymer network and interstitial fluid. The total stress within the gel includes contributions from both the network elasticity and the fluid pressure. The pressure difference across the pillar wall is given by:

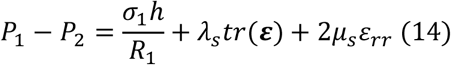

Here, *λ*_s_ and *μ*_s_ are the Lamé coefficients of the gel network, ***ε*** is the strain tensor, and *ε*_rr_ is the radial strain component. Assuming isotropic radial deformation, the radial strain is approximated as: 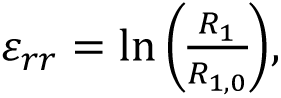, where *R*_1,O_ is the reference (stress-free) pillar radius. Notably, for the parameter values listed in Table 1, Eqs. (13)–(14) predict *P*_2_ − *P*_O_ > 0 and *P*_l_− *P*_2_> 0, corresponding to the pressure ordering *P*_l_> *P*_2_> *P*_O_. At steady state, when all water fluxes vanish, the osmotic pressures follow the same ordering, Π_l_ > Π_2_ > Π_O_.

### Nondimensionalization

We nondimensionalize the governing equations using the characteristic scales of the system. Specifically, we choose the lumen elastic modulus *E*_2_, the initial vesicle radius *R*_2,O_, and the hydraulic time scale *R*_2,O_/(*α*_2_*E*_2_)as the reference stress, length, and time, respectively. The dimensionless system can be written as

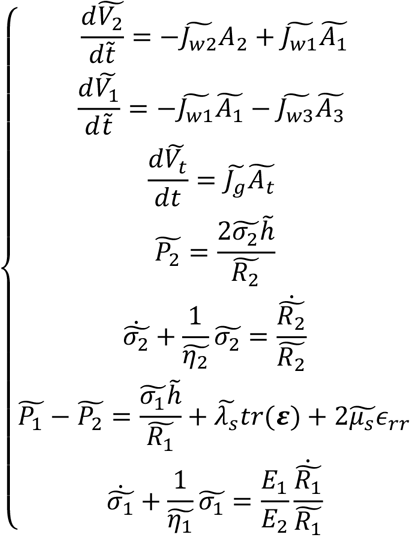

The dimensionless variables are defined as: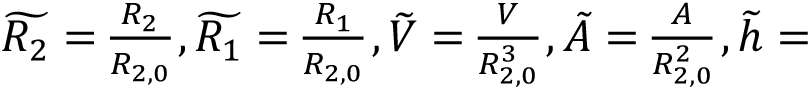 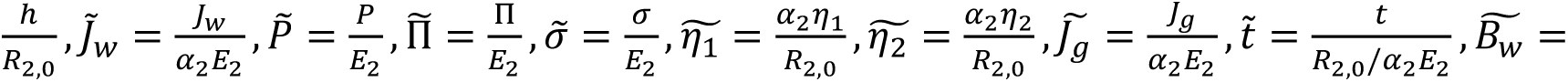 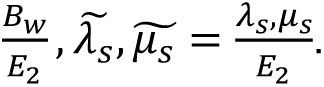 For ion related quantities, we have: 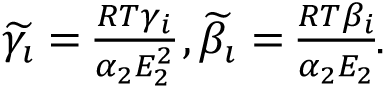. We use: stress scale - *E*, length scale - *R*, time scale – 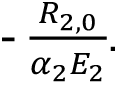. These choices reflect the balance between hydraulic transport and tissue elasticity that governs the system dynamics.

## 2. SIMULATION SETUP

Simulations are performed by iteratively updating transport, geometry, and mechanics in time. At each time step, the following procedure is applied:

### a) Apply perturbations

External perturbations, such as laser ablation or pharmacological treatment, are implemented by modifying the corresponding model parameters or boundary conditions.

For the laser ablation case, we assume that ablation creates a hole in the otic vesicle wall through which luminal fluid escapes via pressure-driven pipe flow. The volumetric outflow is given by *Q*_p_ = (*P*_2_ − *P*_O_)/*ξ*, where *ξ* is the effective hydraulic resistance of the opening. The resistance increases with fluid viscosity and hole depth, and decreases with pore size. Here, *P*_2_ − *P*_O_ is the hydraulic pressure difference between the lumen and the external medium. Because laser ablation experiments probe short timescales (approximately 30 min), we assume that ion transport is negligible and therefore set all ion fluxes to zero. For long-timescale growth dynamics, we assume active ion transport across both the pillar–lumen interface and the lumen wall, corresponding to net ion transport from the pillar into the lumen and from the periotic space into the lumen (*γ*_l_ > 0, *γ*_2_ < 0). Forskolin treatment is modeled as an increase in the overall active ion influx into the lumen, thereby enhancing osmotic water uptake and lumen expansion.

### b) Compute transport fluxes

Using the current geometry, hydraulic pressures, and osmotic pressures, we compute water fluxes and ion fluxes across all relevant interfaces.

### c) Update compartment volumes and ion content

The compartment volumes {*V*_i_} and ion concentrations {*c*_i_} are then updated according to the governing transport equations, Eqs. (4)–(10).

### d) Update geometry

The geometrical variables, including *R*_l_, *R*_2_, and ℎ, are determined from the updated compartment volumes using the geometrical relations in Eqs. (1)–(3). The updated compartment volumes are projected onto the assumed spherical-vesicle and axisymmetric pillar geometries through Eqs. (1)–(3).

### e) Update mechanical state

The viscoelastic stresses and hydraulic pressures are updated using the constitutive relations and pressure–stress balance, Eqs. (11)–(14).

These steps are repeated sequentially until the desired simulation time is reached.

**Table S1.** Model parameters and reference values.

| Parameter | Symbol | Value | Unit | Source |
| --- | --- | --- | --- | --- |
| **Geometrical parameters** |  |  |  |  |
| Reference radius of lumen | $R_{2,0}$ | 46.1 | $\mu\text{m}$ | Estimated from biological data |
| Reference radius of pillar | $R_{1,0}$ | 8.25 | $\mu\text{m}$ | Estimated from biological data |
| Tissue thickness | $h$ | 3.5 | $\mu\text{m}$ | Estimated from biological data |
| **Mechanical parameters** |  |  |  |  |
| Lumen wall elastic modulus | $E_2$ | $10^4$ | Pa | Literature<sup>99,100</sup> |
| Pillar wall elastic modulus | $E_1$ | $10^4$ | Pa | Literature<sup>99,100</sup> |
| Lumen wall viscosity | $\eta_2$ | $3.44 \times 10^7$ | $\text{Pa}\cdot\text{s}$ | Literature<sup>13</sup> |
| Pillar wall viscosity | $\eta_1$ | $2.75 \times 10^6$ | $\text{Pa}\cdot\text{s}$ | Tuned value based on ref<sup>13</sup> |
| Gel Lamé coefficient (bulk) | $\lambda_s$ | 500 | Pa | Assumed |
| Gel Lamé coefficient (shear) | $\mu_s$ | 500 | Pa | Assumed |
| **Transport parameters (water)** |  |  |  |  |
| Lumen wall water permeability | $\alpha_2$ | $7.35 \times 10^{-10}$ | $\text{m}\cdot\text{s}^{-1}\cdot\text{Pa}^{-1}$ | Literature<sup>101</sup> |
| Pillar–lumen permeability | $\alpha_1$ | $7.35 \times 10^{-10}$ | $\text{m}\cdot\text{s}^{-1}\cdot\text{Pa}^{-1}$ | Literature<sup>101</sup> |
| Pillar–external permeability | $\alpha_3$ | $7.35 \times 10^{-10}$ | $\text{m}\cdot\text{s}^{-1}\cdot\text{Pa}^{-1}$ | Literature<sup>101</sup> |
| Effective hydraulic resistance of the ablation opening | $\xi$ | $3.53 \times 10^{13}$ | $\text{Pa}\cdot\text{s}/\text{m}^3$ | Fitted |
| **Transport parameters (ions)** |  |  |  |  |
| Passive ion permeability (pillar–lumen) | $\beta_1$ | $10^{-6} - 10^{-5}$ | $\text{m}\cdot\text{s}^{-1}$ | Literature<sup>101</sup> |
| Passive ion permeability (lumen–external) | $\beta_2$ | $10^{-6} - 10^{-5}$ | $\text{m}\cdot\text{s}^{-1}$ | Literature<sup>101</sup> |
| Active ion pumping (pillar) | $\gamma_1$ | $1.32 \times 10^{-7}$ | $\text{mol}\cdot\text{m}^{-2}\cdot\text{s}^{-1}$ | Literature<sup>101</sup> |
| Active ion pumping (lumen wall) | $\gamma_2$ | $-1.32 \times 10^{-7}$ | $\text{mol}\cdot\text{m}^{-2}\cdot\text{s}^{-1}$ | Literature<sup>101</sup> |
| Fixed charge density (gel) | $c_F$ | 10 | mM | Literature<sup>26,102</sup> |
| External ion concentration | $c_0$ | 323 | mM | Literature<sup>103</sup> |

| Reference scales |  |  |  |  |
| --- | --- | --- | --- | --- |
| Reference length | $R_{2,0}$ | 46.1 | $\mu\text{m}$ | / |
| Reference stress | $E_2$ | $10^4$ | Pa | / |
| Reference time | $t_r$<br>$= R_r$<br>$/(\alpha_2 E_2)$ | 6.27 | s | / |

All scripts used in the theoretical model can be found on GitHub (https://github.com/Yufei-star/Otic_Vesicle_Fluid_Transport_Model).

## Statistics

Plots display individual data points, number of replicates, mean values, and error bars representing standard deviation (SD). Normality of each dataset was assessed using both the Kolmogorov–Smirnov test and the Shapiro–Wilk test prior to parametric statistical analysis. Comparisons between two groups were performed using unpaired or paired two-tailed Student’s t-tests, as appropriate. For datasets that did not follow a normal distribution, the Mann–Whitney U test was used. For multiple comparisons, one-way ANOVA with Tukey’s test for normally distributed data and Kruskal-Wallis test with Dunn’s test for non-normally distributed data were performed. To compute a correlation between normalized change in volume and angle from timelapses, a Spearman rank-order correlation (Spearman’s ρ) was performed on ranked values, with p-value computed from the asymptotic t-distribution for pooled datasets (due to ties in ranks) and an exact p-value computed for individual timelapses. Spearman’s non-parametric approach was used because the data was not assumed to have a linear relationship, due to the measured canalization of pillar angle, beyond which OV lumen volume continues to grow. Statistical analyses and data visualization were performed using GraphPad Prism (versions 9 and 10) and R (version 4.6.0). All analysis scripts used for plotting and statistics in R can be found on GitHub. (https://github.com/MunjalLab/wls_general_analysis).

## Manuscript preparation

The authors used ChatGPT-5 and Claude Sonnet 5 (Anthropic 2026) to improve the clarity and readability of the manuscript text. The authors reviewed and edited the output as needed and take full responsibility for the content of the published article.

## Author Contributions

**Authors:** JW, KH, YW, AB, KL, NS, SS, RH, MB, AM

JW, KH, YW, MB, and AM conceived the project. AM acquired funding, supervised the study, and administered the project. JW performed the majority of the experiments, developed the computational analysis pipeline, and carried out the analyses for all figures, except for the time-lapse imaging, hyaluronidase perturbation and analysis, pillar angle measurements and analysis, and behavioral experiments and MCAM analyses. KH independently acquired the time-lapse imaging datasets, performed the timecourse experiment for Fig. 1 and acquired the HA staining images for Fig. 2 together with JW, performed the experiments and analyses for Fig. 3 together with JW, and contributed to the experiments and analyses for Fig. 5 together with ABB and SS. YW developed and iterated the theoretical model under the supervision of AM and MB. ABB performed the majority of the behavioral experiments with assistance from KL and KH. KL modified and maintained the MCAM system under the supervision of RH and assisted ABB with behavioral data acquisition and analysis under the supervision of MB. NES performed the otic vesicle injections and dissections for single-cell RNA sequencing. JW, KH, and YW prepared the figures with input from AM. JW, KH, YW, and AM interpreted the data and wrote the original manuscript with input from ABB and SS. All authors reviewed and edited the manuscript.

## By Category

Conceptualization: JW, KH, YW, MB, AM

Methodology: JW, KH, YW, AB, KL, RH

Software: JW, KH, YW, AB, KL, SS, RH

Validation: JW, KH, YW, AB, KL, RH, MB, AM

Formal analysis: JW, KH, YW, AB, KL, SS

Investigation: JW, KH, AB, NS

Resources: RH, MB, AM

Data Curation: JW, KH, YW, AB, SS

Writing – Original Draft: JW, KH, YW, AB, KL, SS, AM

Writing – Review & Editing: JW, KH, YW, AB, KL, NS, SS, RH, MB, AM (all authors)

Visualization: JW, KH, YW, AB, KL, SS, AM

Supervision: RH, MB, AM Project administration: AM

Funding acquisition: KH, RH, MB, AM

## Definitions

https://www.elsevier.com/researcher/author/policies-and-guidelines/credit-author-statement

## Acknowledgements

We are thankful to Yusuke Mori and all members of the Principles of Tissue Morphogenesis (Munjal) Lab for their feedback and support. We thank Bagnat and Di Talia Labs for useful discussions. We thank Stefano Di Talia, Ziqi (Alvin) Lu, and Jingwen Shen for feedback on the manuscript. We thank David Schoppik for suggesting uninflated swimbladder control for the *wls* mutants. We thank Eric Liao, Shannon Carroll, and Lisa Tsay for kindly sharing the *wls* mutant and genotyping method. We thank the Duke Zebrafish Core Facility of the Duke School of Medicine for their support. Research reported in this publication was supported by Office of Research Infrastructure Programs (ORIP), the Office of the Director, National Institutes of Health (R44OD024879 and R44OD036187), the National Science Foundation (2036439), and a Duke-Coulter Translational Partnership Grant to RH; National Institute of General Medical Sciences (GM150666) to MB; and by the National Institute of Child Health and Human Development to K.H. (F32HD117588) and to AM (DP2HD115157).

---

**Supplementary Figure 1.**
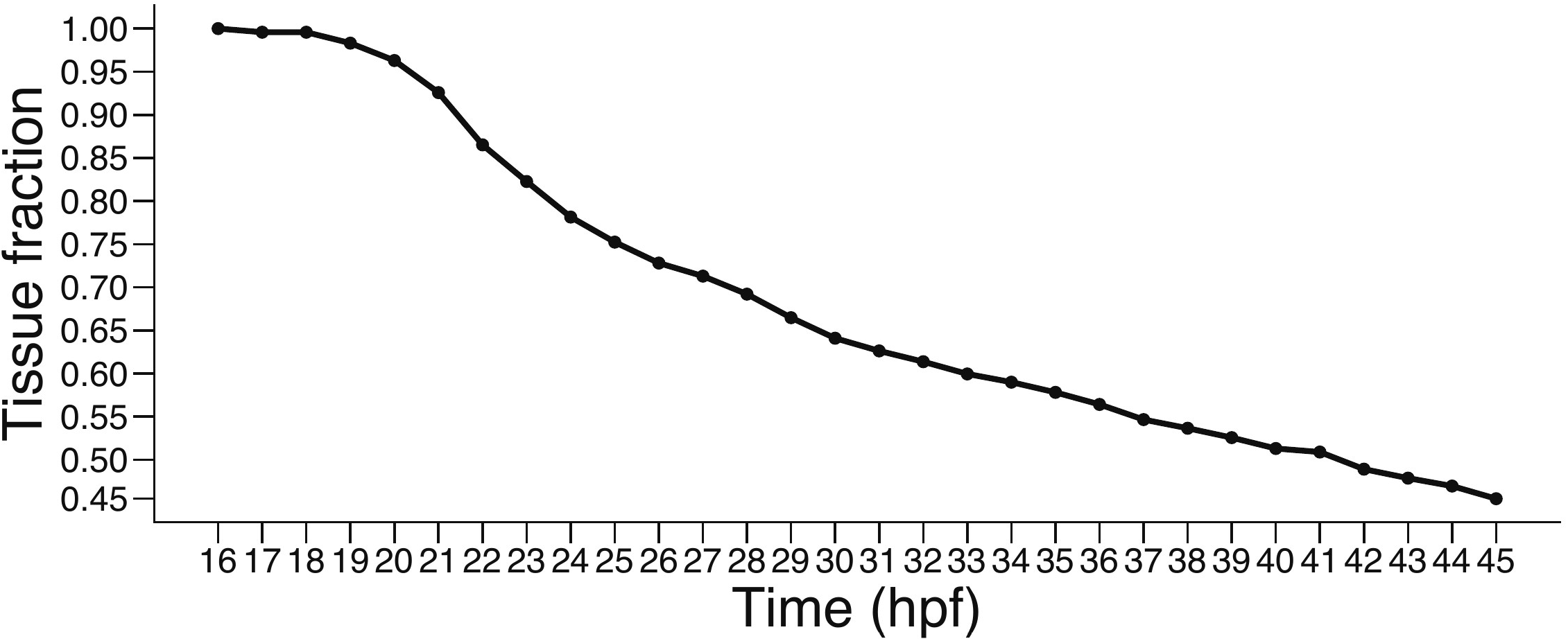
Tissue fraction of OV from early lumen formation to start of morphogenesis, computed from published values, related to. Figure 1. Tissue fraction as a function of tissue volume divided by whole OV volume from 16 to 45 hpf. Data obtained from Fig. 1F in Mosaliganti et al., 2019. Values extracted from figure by Claude AI using corresponding author’s permission.

**Supplementary Figure 2.**
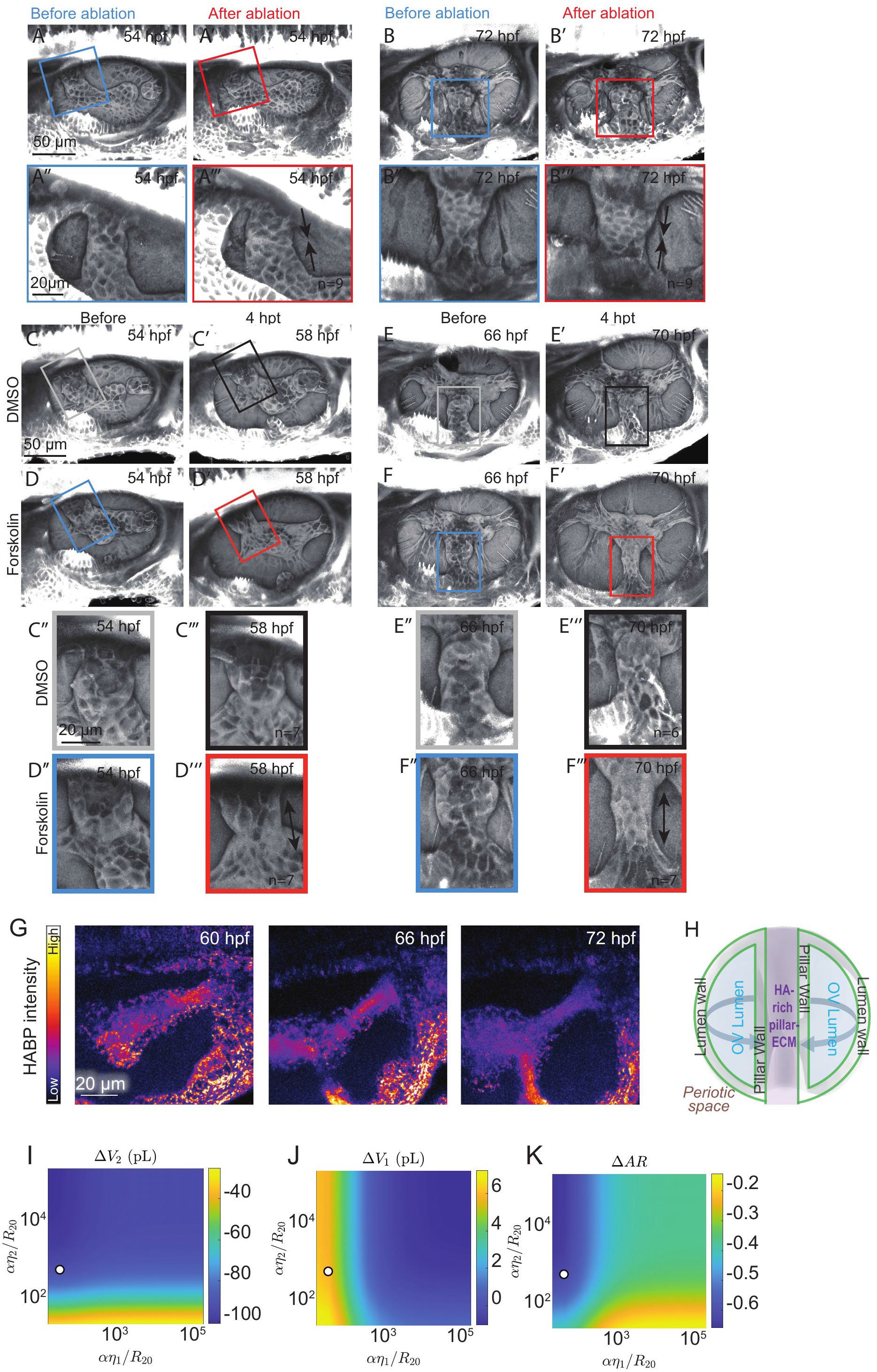
Mechanical coupling of otic vesicle expansion and early pillar morphology, related to. Figure 2**. A-A’.** Representative images of an OV at 54 hpf before and after ablation. **A’’-A’’’**. Magnified insets of the boxed regions in A-A’ showing relaxation of anterior pillar and decreased aspect ratio. **B-B’.** Representative images of an OV at 72 hpf before and after ablation. **B’’-B’’’**. Magnified insets of the boxed regions in C-C’ showing relaxation of ventral pillar and decreased aspect ratio. **C-C’.** Representative images of an OV before and 4 hours post treatment (hpt) by soaking in DMSO at the developmental stages indicated. **C’’-C’’’.** Magnified insets of the boxed regions in C-C’ showing minimal change in anterior pillar aspect ratio after DMSO treatment at the developmental stages indicated. **D-D’.** Representative images of an OV before and 4 hours post treatment (hpt) by soaking in Forskolin solution at the developmental stages indicated. **D’’-D’’’.** Magnified insets of the boxed regions in D-D’ showing elongation of the anterior pillar and increase in aspect ratio after Forskolin treatment at the developmental stages indicated. **E-E’.** Representative images of an OV before and 4 hpt by soaking in DMSO solution at the developmental stages indicated. E’’-E’’’. Magnified insets of the boxed regions in **E-E’** showing elongation of the ventral pillar and modest increase in aspect ratio after DMSO treatment at the developmental stages indicated. **F-F’.** Representative images of an OV before and 4 hpt by soaking in Forskolin solution at the developmental stages indicated. **F’’-F’’’**. Magnified insets of the boxed regions in F-F’ showing elongation of the ventral pillar and increase in aspect ratio after Forskolin treatment at the developmental stages indicated. G. Representative images of the posterior pillars with hyaluronan-binding protein (HABP) staining at the developmental stages indicated. H. Schematic of the otic vesicle structure used in the physical model. **I-K.** Computed phase diagrams showing the changes in lumen volume (Δ*V*_2_), pillar volume (Δ*V*_l_), and pillar aspect ratio (Δ*AR*) over a 30-min period following simulated OV ablation as a function of lumen (2) and pillar (1) epithelium viscosity (*αη*_2_/*R*_2O_, *αη*_l_/*R*_2O_). Here *R*_2O_ denotes the reference lumen radius used for nondimensionalization. The dot indicates the parameter values corresponding to the physiological condition. “n” denotes number of embryos.

**Supplementary Figure 3.**
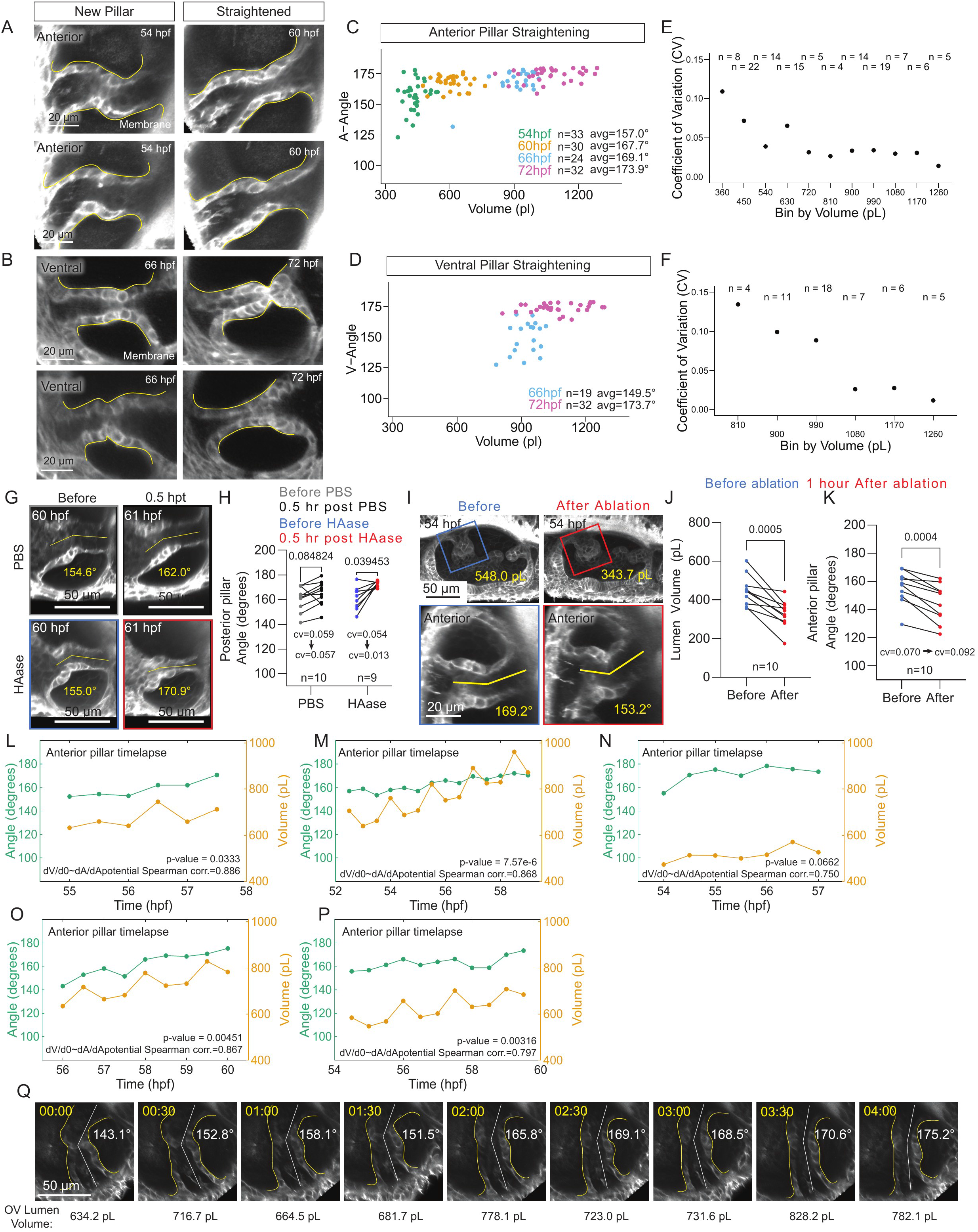
Canalization of pillar morphology via otic vesicle expansion, related to. Figure 3**. A.** Representative confocal images of 3D-rendered anterior pillars from membrane-mNeonGreen-expressing embryos at the stages indicated, demonstrating straightening of anterior pillars after initial shape variability. **B.** Representative confocal images of 3D-rendered ventral pillars from membrane-mNeonGreen-expressing embryos at the stages indicated, demonstrating straightening of ventral pillars after initial shape variability. **C.** Scatter plot of canalization of anterior pillar angle (degrees) vs otic vesicle lumen volume (pL) quantified in embryos pooled at the following developmental stages: 54 hpf (green), 60 hpf (orange), 66 hpf (blue), and 72 hpf (pink). **D.** Scatter plot of canalization of ventral pillar angle (degrees) vs otic vesicle lumen volume (pL) quantified in embryos pooled at the following developmental stages: 66 hpf (blue) and 72 hpf (pink). **E**. Plot of coefficient of variation calculated from embryos in B binned by lumen volume in 90-pL bins. **F.** Plot of coefficient of variation calculated from embryos in E binned by lumen volume in 90-pL bins. **G.** Representative confocal images of 3D-rendered posterior pillars from membrane-mNeonGreen-expressing embryos at the stages indicated, before and 1 hour post treatment (hpt) by periotic injection with 1xPBS as a control or HAase in 1xPBS. **H.** Quantification of posterior pillar angle (degrees) before and 1 hpt periotic injection with 1xPBS or HAase in 1xPBS at 60 hpf. P-values as labeled (two-tailed paired t-test). **I.** Top panel: Representative confocal images of 3D-rendered OVs from membrane-mNeonGreen-expressing embryos at the stages indicated, before and after 2-photon laser ablation of the OV to reduce lumen volume and pressure. Bottom panel: Representative confocal images of 3D-rendered anterior pillars from membrane-mNeonGreen-expressing embryos at the stages indicated, before and after 2-photon laser ablation of the OV to reduce lumen volume and pressure. **J.** Quantification of OV lumen volume (pL) before (blue) and 1 hour after 2-photon laser ablation (red) of the OV at 54 hpf. P-values as labeled (two-tailed paired t-test). **K.** Quantification of anterior pillar angle (degrees) before (blue) and 1 hour after 2-photon laser ablation (red) of the OV at 54 hpf. P-values as labeled (two-tailed paired t-test). **L.** Traces of anterior pillar angle (degrees, green) and OV lumen volume (pL, orange) over time (hours post fertilization, hpf) quantified from one example embryo that was timelapse imaged every 30 minutes from pillar formation to pillar straightening. Spearman correlation coefficient between scaled change in pillar angle and scaled change in lumen volume shown (0.886). **M.** Traces of anterior pillar angle (degrees, green) and OV lumen volume (pL, orange) over time (hpf) quantified from one example embryo that was timelapse imaged every 30 minutes from pillar formation to pillar straightening. Spearman correlation coefficient between change in pillar angle and change in lumen volume shown (0.868). **N.** Traces of anterior pillar angle (degrees, green) and OV lumen volume (pL, orange) over time (hpf) quantified from one example embryo that was timelapse imaged every 30 minutes from pillar formation to pillar straightening. Spearman correlation coefficient between change in pillar angle and change in lumen volume shown (0.750). **O.** Traces of anterior pillar angle (degrees, green) and OV lumen volume (pL, orange) over time (hpf) quantified from one example embryo that was timelapse imaged every 30 minutes from pillar formation to pillar straightening. Spearman correlation coefficient between change in pillar angle and change in lumen volume shown (0.867). **P.** Traces of anterior pillar angle (degrees, green) and OV lumen volume (pL, orange) over time (hpf) quantified from one example embryo that was timelapse imaged every 30 minutes from pillar formation to pillar straightening. Spearman correlation coefficient between change in pillar angle and change in lumen volume shown (0.797). **Q.** Representative time series of anterior pillar cross-sections from one membrane-mNeonGreen-expressing embryo timelapse. Images are each 30 minutes apart, starting from pillar formation. Pillar angle given in each image, and OV lumen volume given below for each timepoint. Pillars outlined in yellow, with angle emphasized with a white line. Coefficient of variation (cv) indicated in certain graphs. Scale bar values indicated in each image.

**Supplementary Figure 4.**
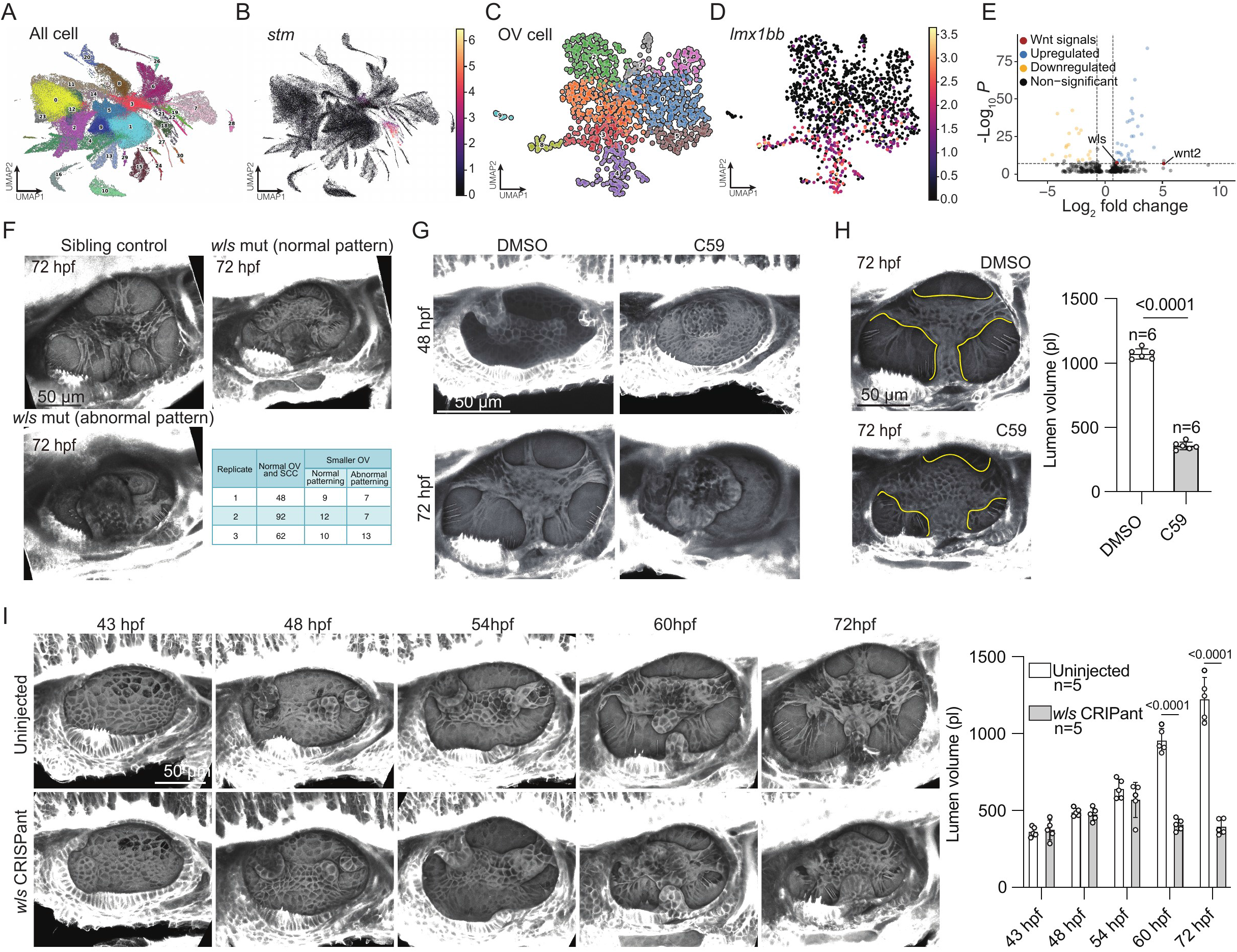
OV expansion is regulated by Wnt signaling through maintenance of lumen barrier function during canal morphogenesis, related to Figure 4. **A.** UMAP dimensionality reduction and Leiden clustering of OV cells in the published inner ear–specific single-cell dataset. **B.** All cells on the UMAP graph with expression of the OV marker *stm* overlain as a heatmap. **C.** All cells in the *stm^+^* OV clusters, colored by subcluster. **D.** All cells in the *stm^+^*OV clusters, with expression of the dorsal OV marker *lmx1bb* overlain as a heatmap, highlighting the pillar-forming cell clusters. **E.** Volcano plot showing differential gene expression between dorsal OV cells and all other OV cells. Genes meeting the significance criteria (adjusted P < 10^-7^ and log2 fold change > 0.7, blue; log2 fold change < −0.7, yellow) are colored by direction of change; non-significant genes are shown in black. Several genes related to Wnt signaling are highlighted (red), including *wntless* (*wls*). **F.** Representative images of 3D-rendered OVs of sibling control (upper-left), *wls* mutant with normal SCC patterning (upper-right), and *wls* mutant with abnormal SCC patterning (lower-left). Lower-right panel: Quantification of otic vesicle phenotypes across biological replicates. **G.** Upper panel: Representative images of 3D-rendered OVs from membrane-mNeonGreen-expressing embryos at 48 hpf after soaking for 24 hours in either DMSO or the Wnt inhibitor C59. Lower panel: Representative images of 3D-rendered OVs from membrane-mNeonGreen-expressing embryos at 72 hpf after DMSO or the Wnt inhibitor C59 treatment from 24-48 hpf. Embryos were washed by egg water at 48 hpf. **H.** Representative images of 3D-rendered OVs from membrane-mNeonGreen-expressing embryos at 72 hpf after soaking for 24 hours in either DMSO or the Wnt inhibitor C59, with pillar region outlined in yellow. Right-hand plot: quantification of OV lumen volume (pL) after 24 hrs of soaking in either DMSO (white) or C59 (grey). **I.** Timecourse imaging of uninjected control embryos (upper panel), and *wls* CRISPants (lower panel). Right-hand plot: quantification of OV lumen volume (pL) of both conditions in the timecourse. “n” denotes the number of embryos. Data are from 1 experiment. Scale bar: 50 μm. Data are mean±s.d. P-values as labeled (unpaired, two-tailed Student’s t-test).

**Supplementary Figure 5.**
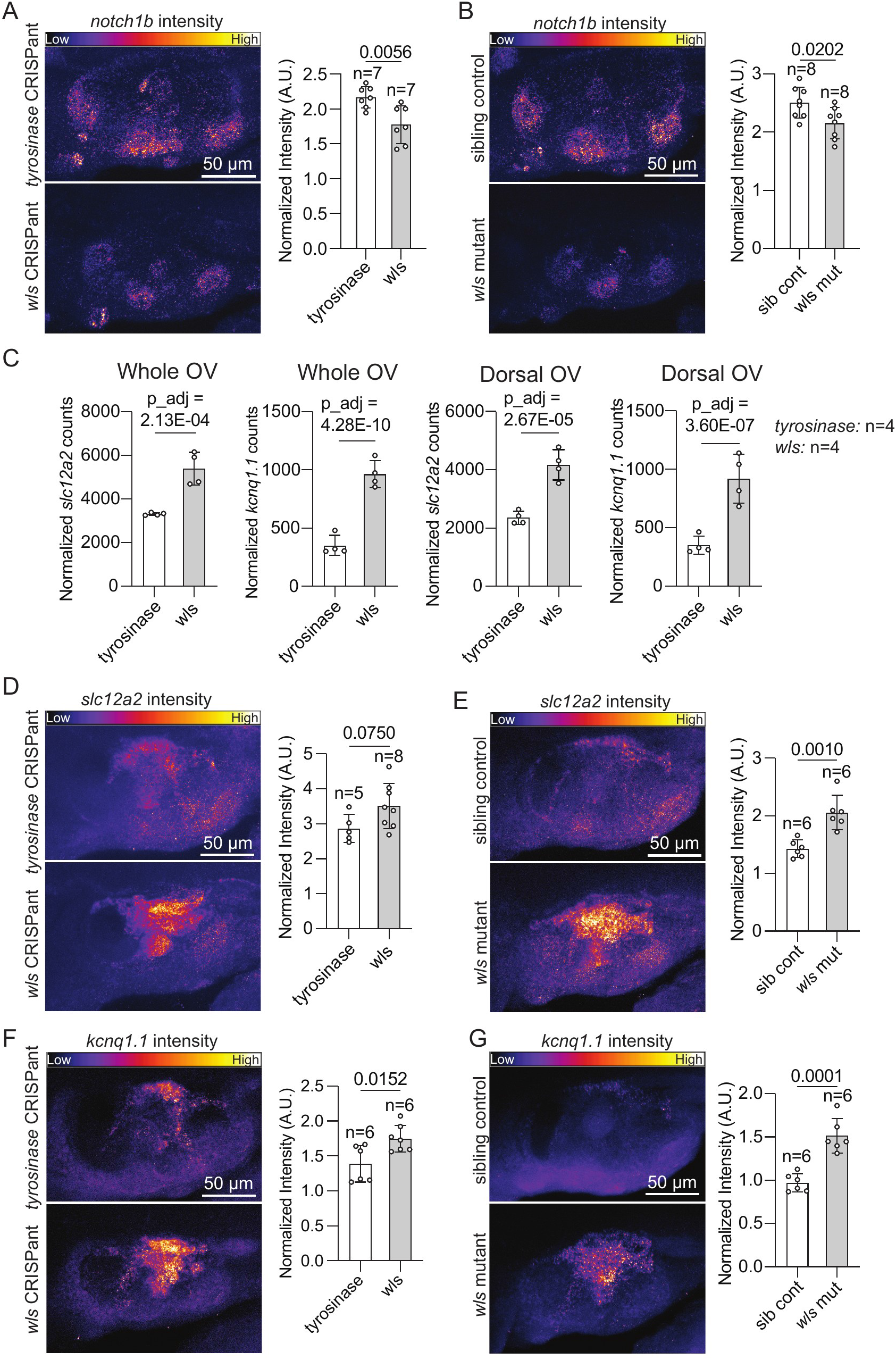
HCR-based validation of Wnt Signaling targets *in vivo* identified by scRNAseq, related to. Figure 4**. A-B.** HCR-FISH on *notch1b* expression in OVs of *wls* and tyrosinase CRISPants (A), sibling controls and *wls* mutants (B). Maximum intensity projections of OVs and quantifications of probe fluorescence intensity are shown. **C.** Pseudobulk analyses of *slc12a2 and kcnq1.1* in cells from *tyrosinase* (white) or *wls* (grey) CRISPants in the whole OV or in the dorsal OV clusters. n denotes the number of biological replicates. Statistical significance was determined using DESeq2, and adjusted P values (Benjamini–Hochberg correction) are indicated. **D-G**. HCR-FISH on *slc12a2* (D, E) *and kcnq1.1* (F, G) expression in OVs of *wls* and tyrosinase CRISPants (D, F), sibling controls and *wls* mutants (E, G). Maximum intensity projections of OVs and quantifications of probe fluorescence intensity are shown. “n” denotes the number of embryos unless otherwise specified. Data are from 1 experiment. Scale bar: 50 μm. Data are mean±s.d. P-values as labeled (unpaired, two-tailed Student’s t-test) unless otherwise specified.

**Supplementary Figure 6.**
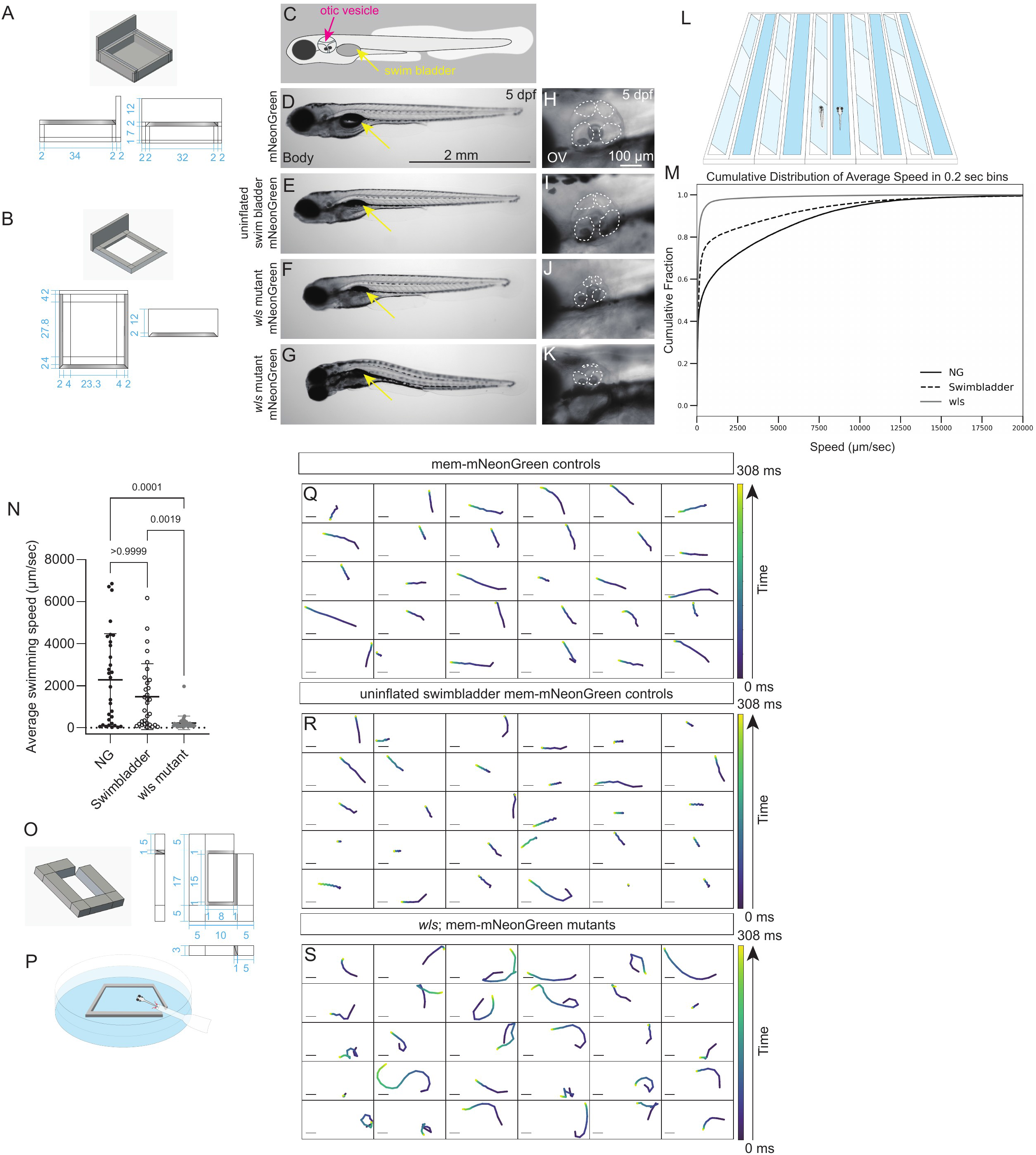
Behavioral analysis of vestibular function upon disruption of Wnt signaling, related to. Figure 5**. A-B.** Schematic illustration of holding chambers used to generate swim bladder-uninflated controls, dimensions given in millimeters (mm). **C.** Schematic illustration of a 5 dpf zebrafish larva, with the swim bladder highlighted in yellow and the otic vesicle highlighted in red. **D-K.** Bright field images of a membrane-mNeonGreen control (D), swim bladder-uninflated control (E), and two *wls* mutants (F-G). Bright field images of the OVs in a membrane-mNeonGreen control (H), swim bladder-uninflated control (I), and two *wls* mutants (J-K). **L.** Schematic illustration of MCAM used to record zebrafish larval swimming behaviors. **M.** Cumulative distribution of average speed of the binned data from membrane-mNeonGreen controls, swim bladder-uninflated controls, and *wls* mutants. **N.** Quantification of average swimming speed (µm/sec) per fish in NG controls, swimbladder controls, or *wls* mutants. Average swimming speed computed from 10 minutes of filmed swimming in a channel (L) under the MCAM. **O.** Schematic illustration of the customized container used in the larval startle response assay, dimensions given in mm. **P.** Schematic illustration showing zebrafish startle response elicited by a directional water current delivered via pipette. **Q-S.** 30 representative swimming paths post startling of membrane-mNeonGreen control (Q), swim bladder-uninflated control (R), and *wls* mutants (S). Scale bars are all 2 mm unless otherwise indicated. Data are mean±s.d. P-values as labeled (Kruskal-Wallis test).

## Video Legends

**Supplementary Video 1. U-Net–based segmentation of the zebrafish otic vesicle lumen, related to Figure 1.**

Representative confocal image stacks of otic vesicles at 40, 48, 54, 60, and 72 hpf are shown with the corresponding U-Net segmentation masks (blue), demonstrating accurate lumen segmentation across developmental stages.

**Supplementary Video 2. Swimming behavior of NG, swimbladder uninflated control, and *wls* mutant fish under normal conditions, related to Figure 5.**

Representative videos of swimming behavior in NG, swimbladder uninflated controls, and *wls* mutants, under normal conditions filmed by the MCAM. Scale bar = 2 mm.

**Supplementary Video 3. Swimming behavior of NG, swimbladder uninflated control, and *wls* mutant fish upon flow-evoked startle response, related to Figure 5.**

Representative videos of swimming behavior in NG, swimbladder uninflated controls, and *wls* mutants, upon flow-evoked startle response. Scale bars = 2 mm.

## References

1. Zhao, X., Tan, R.-S., Tang, H.-C., Teo, S.-K., Su, Y., Wan, M., Leng, S., Zhang, J.-M., Allen, J., Kassab, G.S., et al. (2018). Left Ventricular Wall Stress Is Sensitive Marker of Hypertrophic Cardiomyopathy With Preserved Ejection Fraction. Front. Physiol. 9, 250. 10.3389/fphys.2018.00250.

2. Weibel, E.R. (1970) Morphometric Estimation of Pulmonary Diffusion Capacity. Respir. Physiol. 11, 54–75. 10.1016/0034-5687(70)90102-7.

3. Knust, J., Ochs, M., Gundersen, H.J.G., and Nyengaard, J.R. (2009). Stereological Estimates of Alveolar Number and Size and Capillary Length and Surface Area in Mice Lungs. Anat. Rec. 292, 113–122. 10.1002/ar.20747.

4. Gehr, P., Bachofen, M., and Weibel, E.R. (1978). The normal human lung: ultrastructure and morphometric estimation of diffusion capacity. Respir. Physiol. 32, 121–140. 10.1016/0034-5687(78)90104-4.

5. Wang, J., Wu, Y., Zhou, T., Feng, Y., and Li, L. (2025). Common factors and nutrients affecting intestinal villus height-A review. Anim. Biosci. 38, 1557–1569. 10.5713/ab.25.0002.

6. Garic, M., Vernekar, R., Martín, D.I.Y., Tanguy, S., Loubens, C. de, and Loverdo, C. (2025). Intestinal villi and crypts density maximizing nutrient absorption. Preprint at arXiv, 10.48550/arXiv.2507.03472.

7. Rabbitt, R.D. (2019). Semicircular canal biomechanics in health and disease. J. Neurophysiol. 121, 732–755. 10.1152/jn.00708.2018.

8. Groves, A.K., and Fekete, D.M. (2012). Shaping sound in space: the regulation of inner ear patterning. Development 139, 245–257. 10.1242/dev.067074.

9. Jones, G.M., and Spells, K.E. (1963). A theoretical and comparative study of the functional dependence of the semicircular canal upon its physical dimensions. Proc. R. Soc. Lond. B Biol. Sci. 157, 403–419. 10.1098/rspb.1963.0019.

10. Essner, R.L., Pereira, R.E.E., Blackburn, D.C., Singh, A.L., Stanley, E.L., Moura, M.O., Confetti, A.E., and Pie, M.R. (2022). Semicircular canal size constrains vestibular function in miniaturized frogs. Sci. Adv. 8, eabn1104. 10.1126/sciadv.abn1104.

11. Alsina, B., and Whitfield, T.T. (2017). Sculpting the labyrinth: Morphogenesis of the developing inner ear. Semin. Cell Dev. Biol. 65, 47–59. 10.1016/j.semcdb.2016.09.015.

12. Whitfield, T.T., Riley, B.B., Chiang, M.-Y., and Phillips, B. (2002). Development of the zebrafish inner ear. Dev. Dyn. 223, 427–458. 10.1002/dvdy.10073.

13. Mosaliganti, K.R., Swinburne, I.A., Chan, C.U., Obholzer, N.D., Green, A.A., Tanksale, S., Mahadevan, L., and Megason, S.G. (2019). Size control of the inner ear via hydraulic feedback. eLife 8, e39596. 10.7554/eLife.39596.

14. Munjal, A., Hannezo, E., Tsai, T.Y.-C., Mitchison, T.J., and Megason, S.G. (2021). Extracellular hyaluronate pressure shaped by cellular tethers drives tissue morphogenesis. Cell 184, 6313–6325.e18. 10.1016/j.cell.2021.11.025.

15. Whitfield, T.T. (2025). Chapter Four - Development of the semicircular canals and otolithic organs of the vertebrate inner ear. In Development of Sensory Organs Current Topics in Developmental Biology., G. P. Richardson and D. K. Wu, eds. (Academic Press), pp. 125–184. 10.1016/bs.ctdb.2025.04.001.

16. Mori, Y., Smith, S., Wang, J., Eliora, N., Heikes, K.L., and Munjal, A. (2025). Versican controlled by Lmx1b regulates hyaluronate density and hydration for semicircular canal morphogenesis. Development 152, dev203003. 10.1242/dev.203003.

17. Geng, F.-S., Abbas, L., Baxendale, S., Holdsworth, C.J., Swanson, A.G., Slanchev, K., Hammerschmidt, M., Topczewski, J., and Whitfield, T.T. (2013). Semicircular canal morphogenesis in the zebrafish inner ear requires the function of gpr126 (lauscher), an adhesion class G protein-coupled receptor gene. Dev. Camb. Engl. 140, 4362–4374. 10.1242/dev.098061.

18. Mori, Y., Robin, P., Briggs, A.B., Levic, D.S., Wang, J., Heikes, K.L., Bagnat, M., Hannezo, E., and Munjal, A. (2025). A self-limiting mechanotransduction feedback loop ensures robust organ formation. Preprint at bioRxiv, 10.1101/2025.09.25.678607.

19. Baeza-Loya, S., and Raible, D.W. (2023). Vestibular physiology and function in zebrafish. Front. Cell Dev. Biol. 11, 1172933. 10.3389/fcell.2023.1172933.

20. Hoijman, E., Rubbini, D., Colombelli, J., and Alsina, B. (2015). Mitotic cell rounding and epithelial thinning regulate lumen growth and shape. Nat. Commun. 6, 7355. 10.1038/ncomms8355.

21. Ronneberger, O., Fischer, P., and Brox, T. (2015). U-Net: Convolutional Networks for Biomedical Image Segmentation. In Medical Image Computing and Computer-Assisted Intervention – MICCAI 2015, N. Navab, J. Hornegger, W. M. Wells, and A. F. Frangi, eds. (Springer International Publishing), pp. 234–241. 10.1007/978-3-319-24574-4_28.

22. Dekkers, J.F., Wiegerinck, C.L., de Jonge, H.R., Bronsveld, I., Janssens, H.M., de Winter-de Groot, K.M., Brandsma, A.M., de Jong, N.W.M., Bijvelds, M.J.C., Scholte, B.J., et al. (2013). A functional CFTR assay using primary cystic fibrosis intestinal organoids. Nat. Med. 19, 939–945. 10.1038/nm.3201.

23. Choudhury, M.I., Benson, M.A., and Sun, S.X. (2022). Trans-epithelial fluid flow and mechanics of epithelial morphogenesis. Semin. Cell Dev. Biol. 131, 146–159. 10.1016/j.semcdb.2022.05.020.

24. Torres-Sánchez, A., Kerr Winter, M., and Salbreux, G. (2021). Tissue hydraulics: Physics of lumen formation and interaction. Cells Dev. 168, 203724. 10.1016/j.cdev.2021.203724.

25. Ruiz-Herrero, T., Alessandri, K., Gurchenkov, B.V., Nassoy, P., and Mahadevan, L. (2017). Organ size control via hydraulically gated oscillations. Development 144, 4422–4427. 10.1242/dev.153056.

26. Lai, W.M., Hou, J.S., and Mow, V.C. (1991). A Triphasic Theory for the Swelling and Deformation Behaviors of Articular Cartilage. J. Biomech. Eng. 113, 245–258. 10.1115/1.2894880.

27. Mongera, A., Rowghanian, P., Gustafson, H.J., Shelton, E., Kealhofer, D.A., Carn, E.K., Serwane, F., Lucio, A.A., Giammona, J., and Campàs, O. (2018). A fluid-to-solid jamming transition underlies vertebrate body axis elongation. Nature 561, 401–405. 10.1038/s41586-018-0479-2.

28. Bagnat, M., Daga, B., and Di Talia, S. (2022). Morphogenetic Roles of Hydrostatic Pressure in Animal Development. Annu. Rev. Cell Dev. Biol. 38, 375–394. 10.1146/annurev-cellbio-120320-033250.

29. Chan, C.J., Costanzo, M., Ruiz-Herrero, T., Mönke, G., Petrie, R.J., Bergert, M., Diz-Muñoz, A., Mahadevan, L., and Hiiragi, T. (2019). Hydraulic control of mammalian embryo size and cell fate. Nature 571, 112–116. 10.1038/s41586-019-1309-x.

30. Sur, A., Wang, Y., Capar, P., Margolin, G., Prochaska, M.K., and Farrell, J.A. (2023). Single-cell analysis of shared signatures and transcriptional diversity during zebrafish development. Dev. Cell 58, 3028–3047.e12. 10.1016/j.devcel.2023.11.001.

31. Munjal, A., Kukreja, K., Williams, S., Kawanishi, T., O’Brown, N.M., Ishimatsu, K., Klein, A., Tsung-Megason, S.G., and Swinburne, I.A. (2026). A single-cell transcriptomic atlas of inner ear morphogenesis in zebrafish. Preprint, 10.7554/eLife.111158.1.

32. Bänziger, C., Soldini, D., Schütt, C., Zipperlen, P., Hausmann, G., and Basler, K. (2006). Wntless, a Conserved Membrane Protein Dedicated to the Secretion of Wnt Proteins from Signaling Cells. Cell 125, 509–522. 10.1016/j.cell.2006.02.049.

33. Thisse, B., Pflumio, S., Fürthauer, M., Loppin, B., Heyer, V., Degrave, A., Woehl, R., Lux, A., Steffan, T., Charbonnier, X.Q., et al. (2001). Expression of the zebrafish genome during embryogenesis (NIH R01 RR15402) (ZFIN Direct Data Submission). https://zfin.org/ZDB-PUB-010810-1.

34. Choi, H.M.T., Beck, V.A., and Pierce, N.A. (2014). Next-Generation in Situ Hybridization Chain Reaction: Higher Gain, Lower Cost, Greater Durability. ACS Nano 8, 4284–4294. 10.1021/nn405717p.

35. Kuan, Y.-S., Roberson, S., Akitake, C.M., Fortuno, L., Gamse, J., Moens, C., and Halpern, M.E. (2015). Distinct requirements for Wntless in habenular development. Dev. Biol. 406, 117–128. 10.1016/j.ydbio.2015.06.006.

36. Rochard, L., Monica, S.D., Ling, I.T.C., Kong, Y., Roberson, S., Harland, R., Halpern, M., and Liao, E.C. (2016). Roles of Wnt pathway genes wls, wnt9a, wnt5b, frzb and gpc4 in regulating convergent-extension during zebrafish palate morphogenesis. Development 143, 2541–2547. 10.1242/dev.137000.

37. Proffitt, K.D., Madan, B., Ke, Z., Pendharkar, V., Ding, L., Lee, M.A., Hannoush, R.N., and Virshup, D.M. (2013). Pharmacological Inhibition of the Wnt Acyltransferase PORCN Prevents Growth of WNT-Driven Mammary Cancer. Cancer Res. 73, 502–507. 10.1158/0008-5472.CAN-12-2258.

38. Söllner, C., Burghammer, M., Busch-Nentwich, E., Berger, J., Schwarz, H., Riekel, C., and Nicolson, T. (2003). Control of Crystal Size and Lattice Formation by Starmaker in Otolith Biomineralization. Science 302, 282–286. 10.1126/science.1088443.

39. Żak, M., and Daudet, N. (2021). A gradient of Wnt activity positions the neurosensory domains of the inner ear. eLife 10, e59540. 10.7554/eLife.59540.

40. Jayasena, C.S., Ohyama, T., Segil, N., and Groves, A.K. (2008) Notch signaling augments the canonical Wnt pathway to specify the size of the otic placode. Development 135, 2251–2261. 10.1242/dev.017905.

41. Abbas, L., and Whitfield, T.T. (2009). Nkcc1 (Slc12a2) is required for the regulation of endolymph volume in the otic vesicle and swim bladder volume in the zebrafish larva. Dev. Camb. Engl. 136, 2837–2848. 10.1242/dev.034215.

42. Collinet, C., and Lecuit, T. (2021). Programmed and self-organized flow of information during morphogenesis. Nat. Rev. Mol. Cell Biol. 22, 245–265. 10.1038/s41580-020-00318-6.

43. Chugh, M., Munjal, A., and Megason, S.G. (2022). Hydrostatic pressure as a driver of cell and tissue morphogenesis. Semin. Cell Dev. Biol. 131, 134–145. 10.1016/j.semcdb.2022.04.021.

44. Nusse, R., and Clevers, H. (2017). Wnt/β-Catenin Signaling, Disease, and Emerging Therapeutic Modalities. Cell 169, 985–999. 10.1016/j.cell.2017.05.016.

45. Valenta, T., Hausmann, G., and Basler, K. (2012). The many faces and functions of β-catenin. EMBO J. 31, 2714–2736. 10.1038/emboj.2012.150.

46. Liebner, S., Corada, M., Bangsow, T., Babbage, J., Taddei, A., Czupalla, C.J., Reis, M., Felici, A., Wolburg, H., Fruttiger, M., et al. (2008). Wnt/β-catenin signaling controls development of the blood–brain barrier. J. Cell Biol. 183, 409–417. 10.1083/jcb.200806024.

47. Nicolson, T., Rüsch, A., Friedrich, R.W., Granato, M., Ruppersberg, J.P., and Nüsslein-Volhard, C. (1998). Genetic Analysis of Vertebrate Sensory Hair Cell Mechanosensation: the Zebrafish Circler Mutants. Neuron 20, 271–283. 10.1016/S0896-6273(00)80455-9.

48. Hammond, K.L., Loynes, H.E., Mowbray, C., Runke, G., Hammerschmidt, M., Mullins, M.C., Hildreth, V., Chaudhry, B., and Whitfield, T.T. (2009). A Late Role for bmp2b in the Morphogenesis of Semicircular Canal Ducts in the Zebrafish Inner Ear. PLOS ONE 4, e4368. 10.1371/journal.pone.0004368.

49. Portugues, R., and Engert, F. (2011). Adaptive Locomotor Behavior in Larval Zebrafish. Front. Syst. Neurosci. 5, 72. 10.3389/fnsys.2011.00072.

50. Beppi, C., Straumann, D., and Bögli, S.Y. (2021). A model-based quantification of startle reflex habituation in larval zebrafish. Sci. Rep. 11, 846. 10.1038/s41598-020-79923-6.

51. Wang, C., Zhong, Z., Sun, P., Zhong, H., Li, H., and Chen, F. (2017). Evaluation of the Hair Cell Regeneration in Zebrafish Larvae by Measuring and Quantifying the Startle Responses. Neural Plast. 2017, 8283075. 10.1155/2017/8283075.

52. Kimmel, C.B., Patterson, J., and Kimmel, R.O. (1974). The development and behavioral characteristics of the startle response in the zebra fish. Dev. Psychobiol. 7, 47–60. 10.1002/dev.420070109.

53. Thomson, E.E., Harfouche, M., Kim, K., Konda, P.C., Seitz, C.W., Cooke, C., Xu, S., Jacobs, W.S., Blazing, R., Chen, Y., et al. (2022). Gigapixel imaging with a novel multi-camera array microscope. eLife 11, e74988. 10.7554/eLife.74988.

54. Lindsey, B.W., Smith, F.M., and Croll, R.P. (2010). From Inflation to Flotation: Contribution of the Swimbladder to Whole-Body Density and Swimming Depth During Development of the Zebrafish (Danio rerio). Zebrafish 7, 85–96. 10.1089/zeb.2009.0616.

55. Venuto, A., Thibodeau-Beganny, S., Trapani, J.G., and Erickson, T. (2023). A sensation for inflation: initial swim bladder inflation in larval zebrafish is mediated by the mechanosensory lateral line. J. Exp. Biol. 226, jeb245635. 10.1242/jeb.245635.

56. Goolish, E.M., and Okutake, K. (1999). Lack of gas bladder inflation by the larvae of zebrafish in the absence of an air-water interface. J. Fish Biol. 55, 1054–1063. 10.1111/j.1095-8649.1999.tb00740.x.

57. Abbas, L., and Whitfield, T.T. (2009). Nkcc1 (Slc12a2) is required for the regulation of endolymph volume in the otic vesicle and swim bladder volume in the zebrafish larva. Development 136, 2837–2848. 10.1242/dev.034215.

58. Riley, B.B., and Moorman, S.J. (2000). Development of utricular otoliths, but not saccular otoliths, is necessary for vestibular function and survival in zebrafish. J. Neurobiol. 43, 329–337. 10.1002/1097-4695(20000615)43:4%3C329::AID-NEU2%3E3.0.CO;2-H.

59. Trinh, D.-C., Alonso-Serra, J., Asaoka, M., Colin, L., Cortes, M., Malivert, A., Takatani, S., Zhao, F., Traas, J., Trehin, C., et al. (2021). How Mechanical Forces Shape Plant Organs. Curr. Biol. 31, R143–R159. 10.1016/j.cub.2020.12.001.

60. Beauzamy, L., Nakayama, N., and Boudaoud, A. (2014). Flowers under pressure: ins and outs of turgor regulation in development. Ann. Bot. 114, 1517–1533. 10.1093/aob/mcu187.

61. Naganathan, S.R., Popović, M., and Oates, A.C. (2022). Left–right symmetry of zebrafish embryos requires somite surface tension. Nature 605, 516–521. 10.1038/s41586-022-04646-9.

62. Hong, L., Dumond, M., Tsugawa, S., Sapala, A., Routier-Kierzkowska, A.-L., Zhou, Y., Chen, C., Kiss, A., Zhu, M., Hamant, O., et al. (2016). Variable Cell Growth Yields Reproducible Organ Development through Spatiotemporal Averaging. Dev. Cell 38, 15–32. 10.1016/j.devcel.2016.06.016.

63. Naganathan, S.R. (2024). An emerging role for tissue plasticity in developmental precision. Biochem. Soc. Trans. 52, 987–995. 10.1042/BST20230173.

64. Chan, C.J., and Hiiragi, T. (2020). Integration of luminal pressure and signalling in tissue self-organization. Development 147, dev181297. 10.1242/dev.181297.

65. Hannezo, E., and Heisenberg, C.-P. (2022). Rigidity transitions in development and disease. Trends Cell Biol. 32, 433–444. 10.1016/j.tcb.2021.12.006.

66. Lecuit, T., and Lenne, P.-F. (2007). Cell surface mechanics and the control of cell shape, tissue patterns and morphogenesis. Nat. Rev. Mol. Cell Biol. 8, 633–644. 10.1038/nrm2222.

67. Rozario, T., and DeSimone, D.W. (2010). The extracellular matrix in development and morphogenesis: A dynamic view. Dev. Biol. 341, 126–140. 10.1016/j.ydbio.2009.10.026.

68. Miller, R.K., and McCrea, P.D. (2010). Wnt to build a tube: contributions of Wnt signaling to epithelial tubulogenesis. Dev. Dyn. Off. Publ. Am. Assoc. Anat. 239, 77–93. 10.1002/dvdy.22059.

69. MacDonald, B.T., Tamai, K., and He, X. (2009). Wnt/β-Catenin Signaling: Components, Mechanisms, and Diseases. Dev. Cell 17, 9–26. 10.1016/j.devcel.2009.06.016.

70. Galceran, J., Sustmann, C., Hsu, S.-C., Folberth, S., and Grosschedl, R. (2004). LEF1-mediated regulation of *Delta-like1* links Wnt and Notch signaling in somitogenesis. Genes Dev. 18, 2718–2723. 10.1101/gad.1249504.

71. Najarro, E.H., Huang, J., Jacobo, A., Quiruz, L.A., Grillet, N., and Cheng, A.G. (2020). Dual regulation of planar polarization by secreted Wnts and Vangl2 in the developing mouse cochlea. Development 147, dev191981. 10.1242/dev.191981.

72. Swinburne, I.A., Mosaliganti, K.R., Upadhyayula, S., Liu, T.-L., Hildebrand, D.G.C., Tsai, T.Y.-C., Chen, A., Al-Obeidi, E., Fass, A.K., Malhotra, S., et al. (2018). Lamellar projections in the endolymphatic sac act as a relief valve to regulate inner ear pressure. eLife 7, e37131. 10.7554/eLife.37131.

73. Gürkov, R., Pyykö, I., Zou, J., and Kentala, E. (2016). What is Menière’s disease? A contemporary re-evaluation of endolymphatic hydrops. J. Neurol. 263, 71–81. 10.1007/s00415-015-7930-1.

74. Johns, J.D., Olszewski, R., Strepay, D., Lopez, I.A., Ishiyama, A., and Hoa, M. (2023). Emerging Mechanisms in the Pathogenesis of Menière’s Disease: Evidence for the Involvement of Ion Homeostatic or Blood–Labyrinthine Barrier Dysfunction in Human Temporal Bones. Otol. Neurotol. 44, 1057–1065. 10.1097/MAO.0000000000004016.

75. Levic, D.S., Yamaguchi, N., Wang, S., Knaut, H., and Bagnat, M. (2021). Knock-in tagging in zebrafish facilitated by insertion into non-coding regions. Development 148, dev199994. 10.1242/dev.199994.

76. Ota, S., Hisano, Y., Muraki, M., Hoshijima, K., Dahlem, T.J., Grunwald, D.J., Okada, Y., and Kawahara, A. (2013). Efficient identification of TALEN-mediated genome modifications using heteroduplex mobility assays. Genes Cells Devoted Mol. Cell. Mech. 18, 450–458. 10.1111/gtc.12050.

77. Paszke, A., Gross, S., Massa, F., Lerer, A., Bradbury, J., Chanan, G., Killeen, T., Lin, Z., Gimelshein, N., Antiga, L., et al. (2019) PyTorch: An Imperative Style, High-Performance Deep Learning Library. CoRR, abs/1912.01703. 10.48550/arXiv.1912.01703.

78. Yushkevich, P.A., Piven, J., Hazlett, H.C., Smith, R.G., Ho, S., Gee, J.C., and Gerig, G. (2006). User-guided 3D active contour segmentation of anatomical structures: Significantly improved efficiency and reliability. NeuroImage 31, 1116–1128. 10.1016/j.neuroimage.2006.01.015.

79. Solovyev, R., Kalinin, A.A., and Gabruseva, T. (2022). 3D convolutional neural networks for stalled brain capillary detection. Comput. Biol. Med. 141, 105089. 10.1016/j.compbiomed.2021.105089.

80. He, K., Zhang, X., Ren, S., and Sun, J. (2016). Deep Residual Learning for Image Recognition. In 2016 IEEE Conference on Computer Vision and Pattern Recognition (CVPR), pp. 770–778. 10.1109/CVPR.2016.90.

81. Sorlien, E.L., Witucki, M.A., and Ogas, J. (2018). Efficient Production and Identification of CRISPR/Cas9-generated Gene Knockouts in the Model System Danio rerio. J. Vis. Exp. JoVE, 56969. 10.3791/56969.

82. Swinburne, I.A., Mosaliganti, K.R., Green, A.A., and Megason, S.G. (2015). Improved Long-Term Imaging of Embryos with Genetically Encoded α-Bungarotoxin. PLOS ONE 10, e0134005. 10.1371/journal.pone.0134005.

83. Schindelin, J., Arganda-Carreras, I., Frise, E., Kaynig, V., Longair, M., Pietzsch, T., Preibisch, S., Rueden, C., Saalfeld, S., Schmid, B., et al. (2012). Fiji: an open-source platform for biological-image analysis. Nat. Methods 9, 676–682. 10.1038/nmeth.2019.

84. Golob, J.L., Margolis, E., Hoffman, N.G., and Fredricks, D.N. (2017). Evaluating the accuracy of amplicon-based microbiome computational pipelines on simulated human gut microbial communities. BMC Bioinformatics 18, 283. 10.1186/s12859-017-1690-0.

85. Korostin, D., Kulemin, N., Naumov, V., Belova, V., Kwon, D., and Gorbachev, A. (2020). Comparative analysis of novel MGISEQ-2000 sequencing platform vs Illumina HiSeq 2500 for whole-genome sequencing. PLOS ONE 15, e0230301. 10.1371/journal.pone.0230301.

86. Huang, J., Liang, X., Xuan, Y., Geng, C., Li, Y., Lu, H., Qu, S., Mei, X., Chen, H., Yu, T., et al. (2017). A reference human genome dataset of the BGISEQ-500 sequencer. GigaScience 6, gix024. 10.1093/gigascience/gix024.

87. Farrell, J.A., Wang, Y., Riesenfeld, S.J., Shekhar, K., Regev, A., and Schier, A.F. (2018). Single-cell reconstruction of developmental trajectories during zebrafish embryogenesis. Science 360, eaar3131. 10.1126/science.aar3131.

88. Butler, A., Hoffman, P., Smibert, P., Papalexi, E., and Satija, R. (2018). Integrating single-cell transcriptomic data across different conditions, technologies, and species. Nat. Biotechnol. 36, 411–420. 10.1038/nbt.4096.

89. Hao, Y., Hao, S., Andersen-Nissen, E., Mauck, W.M., Zheng, S., Butler, A., Lee, M.J., Wilk, A.J., Darby, C., Zager, M., et al. (2021). Integrated analysis of multimodal single-cell data. Cell 184, 3573–3587.e29. 10.1016/j.cell.2021.04.048.

90. Stuart, T., Butler, A., Hoffman, P., Hafemeister, C., Papalexi, E., Mauck, W.M., Hao, Y., Stoeckius, M., Smibert, P., and Satija, R. (2019). Comprehensive Integration of Single-Cell Data. Cell 177, 1888–1902.e21. 10.1016/j.cell.2019.05.031.

91. Wolf, F.A., Angerer, P., and Theis, F.J. (2018). SCANPY: large-scale single-cell gene expression data analysis. Genome Biol. 19, 15. 10.1186/s13059-017-1382-0.

92. Howe, K., Clark, M.D., Torroja, C.F., Torrance, J., Berthelot, C., Muffato, M., Collins, J.E., Humphray, S., McLaren, K., Matthews, L., et al. (2013). The zebrafish reference genome sequence and its relationship to the human genome. Nature 496, 498–503. 10.1038/nature12111.

93. Harrison, P.W., Amode, M.R., Austine-Orimoloye, O., Azov, A.G., Barba, M., Barnes, I., Becker, A., Bennett, R., Berry, A., Bhai, J., et al. (2024). Ensembl 2024. Nucleic Acids Res. 52, D891–D899. 10.1093/nar/gkad1049.

94. Love, M.I., Huber, W., and Anders, S. (2014). Moderated estimation of fold change and dispersion for RNA-seq data with DESeq2. Genome Biol. 15, 550. 10.1186/s13059-014-0550-8.

95. Chen, H., Li, K., Kreiss, L., Reamey, P., Pierce, L.X., Zhang, R., Da Luz, R., Chaware, A., Kim, K., Cook, C.B., et al. (2026). High-throughput multi-camera array microscope platform for automated 3D behavioral analysis of swimming zebrafish larvae. Commun. Biol. 9, 141. 10.1038/s42003-025-09421-w.

96. Harfouche, M., Kim, K., Zhou, K.C., Konda, P.C., Sharma, S., Thomson, E.E., Cooke, C., Xu, S., Kreiss, L., Chaware, A., et al. (2023). Imaging across multiple spatial scales with the multi-camera array microscope. Optica 10, 471–480. 10.1364/OPTICA.478010.

97. Girstmair, J., Pietzsch, T., Ulman, V., Hahmann, S., Arzt, M., Handberg-Thorsager, M., Pantze, S., Sugawara, K., Haase, R., Tinevez, J.-Y., et al. (2025). Mastodon: the Command Center for Large-Scale Lineage-Tracing Microscopy Datasets. Preprint at bioRxiv, 10.64898/2025.12.10.693416.

98. Gallois, B., and Candelier, R. (2021). FastTrack: An open-source software for tracking varying numbers of deformable objects. PLOS Comput. Biol. 17, e1008697. 10.1371/journal.pcbi.1008697.

99. Chevalier, N.R., Gazquez, E., Dufour, S., and Fleury, V. (2016). Measuring the micromechanical properties of embryonic tissues. Methods 94, 120–128. 10.1016/j.ymeth.2015.08.001.

100. Kuznetsova, T.G., Starodubtseva, M.N., Yegorenkov, N.I., Chizhik, S.A., and Zhdanov, R.I. (2007). Atomic force microscopy probing of cell elasticity. Micron 38, 824–833. 10.1016/j.micron.2007.06.011.

101. Jiang, H., and Sun, S.X. (2013). Cellular Pressure and Volume Regulation and Implications for Cell Mechanics. Biophys. J. 105, 609–619. 10.1016/j.bpj.2013.06.021.

102. Mow, V.C., Ateshian, G.A., Lai, W.M., and Gu, W.Y. (1998). Effects of fixed charges on the stress–relaxation behavior of hydrated soft tissues in a confined compression problem. Int. J. Solids Struct. 35, 4945–4962. 10.1016/S0020-7683(98)00103-6.

103. Vian, A., Pochitaloff, M., Yen, S.-T., Kim, S., Pollock, J., Liu, Y., Sletten, E.M., and Campàs, O. (2023). In situ quantification of osmotic pressure within living embryonic tissues. Nat. Commun. 14, 7023. 10.1038/s41467-023-42024-9.

